# miR-631 suppresses oncogene *RAB11A* in oral squamous cell carcinoma

**DOI:** 10.1101/2025.04.26.650806

**Authors:** Harsha Pulakkat Muralidharan, Radha Mishra, Champaka Gopal, Arun Kumar

**Author notes:** Joint corresponding authors: Email ID (HPM), Email ID (AK).

## Abstract

Despite technological advancements, the five-year survival rate for oral squamous cell carcinoma (OSCC) remains dismally low. To develop more effective therapies, we intended to identify the potential therapeutic tumor suppressor microRNAs in OSCC. Previously, a microRNA microarray analysis of 5-Azacytidine treated SCC131 cells had identified 50 upregulated miRs. Of these, miR-631 that had no previously known roles in OSCC was selected for further analysis in the present study. We show that miR-631 binds directly to the oncogene *RAB11A* and reduces its transcript and protein levels. The upregulation of miR-631 occurs due to its gene promoter demethylation after 5-Azacytidine treatment. We demonstrate that miR-631 reduces proliferation and anchorage-independent growth of OSCC cells in soft agar and promotes apoptosis, in part, via targeting *RAB11A*. An inverse relationship between miR-631 and *RAB11A* levels observed across multiple cancer cell lines and in 58.33% of OSCC patient samples highlights the biological significance of their interaction. Further, miR-631 and *RAB11A* interaction reduces the Wnt signaling in OSCC. Additionally, the nude mice OSCC xenograft study proves the tumor suppressive nature of miR-631. Based on our results, we propose that miR-631 holds promise as a potential therapeutic agent for treating OSCC.

## Introduction

Oral squamous cell carcinoma (OSCC) accounts for 90% of all oral cancers [1]. The Global Cancer Observatory reported 377,713 cases of OSCC in 2022, with a projected 40% rise by 2040 [GLOBOCAN 2022; https://gco.iarc.fr/]. Despite advancements, its five-year survival rate remains below 50%, underscoring the urgent need for effective therapeutic strategies [2]. OSCC is driven by risk factors including tobacco and alcohol consumption, smoking, betel quid chewing [3], HPV infections [4], nutrient deficiencies [5], genetic susceptibility, and poor oral hygiene [6].

Current treatments for OSCC include combinations of surgery, radiation therapy, chemotherapy, immunotherapy, and targeted approaches [7]. While these methods have improved outcomes to some extent, the search for innovative therapies continues. In this context, microRNAs (miRs) have emerged as promising therapeutic agents. These small, non-coding RNA molecules, typically ∼18-25 nucleotides long, regulate gene expression by binding to complementary sequences in target genes [8]. miRs influence diverse cellular processes, including cell-cell communication, development, immune responses, epigenetic regulation, and hallmarks of cancer [9]. Most commonly, they target the 3’ untranslated regions (3’UTRs) of genes to inhibit translation or to degrade the RNA transcripts. miRs have also been shown to interact with coding regions, 5’UTRs and promoter regions [9, 10, 11].

miRs can act either as oncomiRs [12], which positively regulate oncogenes, or as tumor suppressor miRs [13,14], which negatively regulate oncogenes. Factors such as the amplification or deletion of miR genes, defects in miR biogenesis pathways, alterations in miR transcription control, and dysregulated epigenetic modifications can contribute to cancer development [15]. Many miRs are epigenetically dysregulated in OSCC [16]. Further, OSCC patient tissue samples exhibit higher levels of gene hypermethylation compared to their normal counterparts [17]. Hypermethylation of gene promoters can silence gene expression [10, 14]. Reversing such methylation patterns can reactivate these genes, offering a potential therapeutic strategy [18]. Similarly, tumor suppressor miR genes are also susceptible to hypermethylation in cancer cells. Profiling the methylation status of miRs in OSCC could uncover novel tumor suppressors, paving the way for innovative therapeutic approaches. With this rationale, More et al. [13] have treated OSCC cells with the DNMT inhibitor 5-Azacytidine [13]. Using the microRNA microarray analysis, they have showed the upregulation of 50 miRs and the downregulation of 28 miRs, including the upregulation of miR-631 with a significant fold change of 4.39 [13]. Given the upregulation of miR-631 in the microRNA microarray analysis [13] and since nothing is known about this microRNA in OSCC, we conducted further experiments to investigate the role of miR-631 in OSCC pathogenesis. The present study reports the mechanism behind its upregulation in 5-Azacytidine treated OSCC cells and the impact on various cancer hallmarks. We also report the biological relevance of miR-631 and its target gene *RAB11A* interaction in patient tumor samples, as well as its therapeutic potential in nude mice.

## Results

### Upregulation of miR-631 in 5-Azacytidine treated SCC131 cells

As anticipated, qRT-PCR revealed a significant increase in miR-631 expression in 5-Azacytidine treated SCC131 cells relative to DMSO (control) treated cells, thereby corroborating the findings from the microRNA microarray analysis **(Fig. S1a)**. A similar trend was observed through semi-quantitative RT-PCR **(Fig. S1b)**.

### miR-631 overexpression reduces proliferation of OSCC cells

Since miR-631 level increased in SCC131 cells after the 5-Azacytidine treatment, we aimed to determine if it has a tumor suppressive nature. First, miR-631 was cloned in the pcDNA3-EGFP vector to generate the pmiR-631 construct. This construct and the vector were transiently transfected separately in cells from SCC131 and SCC084 cell lines, and the cell proliferation was assessed using the trypan blue dye exclusion assay. Both SCC131 and SCC084 cells showed reduced proliferation during a period of 4 days upon transfection with pmiR-631 in comparison to those transfected with the vector **(Fig. S2a)**. The effect was further confirmed by the BrdU incorporation assay in cells from both cell lines **(Fig. S2b)**. These findings indicated the anti-proliferative nature of miR-631.

### Demethylation of the *MIR631* promoter causes miR-631 upregulation upon 5-Azacytidine treatment

Since miR-631 was upregulated post 5-Azacytidine treatment, we proposed that this might be due to the removal of methylation from the *MIR631* promoter. Since the promoter for this intronic miR remains uncharacterized, we sought to characterize the promoter of the *MIR631* gene, which is located in an intron between exons 5 and 6 in the antisense strand of the host gene *NEIL1* (Nei like DNA glycosylase 1) on chromosome 15 **(Fig. S3)**. Based on the bioinformatics prediction using DBTSS, Promoter 2.0 and Alggen Promo databases, a putative *MIR631* promoter sequence was retrieved **(Fig. S4)**. We cloned it in the pGL3-basic vector to generate pProm-MIR631. SCC131 cells then underwent transient transfection with the constructs (pProm-MIR631, pGL3-control (positive control) and pGL3-basic (negative control)) separately. In a dual-luciferase reporter assay, the cloned *MIR631* sequence in pProm-MIR631 showed an enhanced luciferase activity as compared to the pGL3-basic, confirming that it represents the *MIR631* promoter (**Fig. 1a**). We then looked for the methylation of a CpG-rich region in the *MIR631* promoter, using the sodium bisulfite PCR sequencing (**Fig. 1b**). The findings revealed 70% methylation of CpG sites following 5-Azacytidine exposure in contrast to 98.60% methylation in control cells (**Fig. 1c**), indicating that reduced methylation at the promoter region may be responsible for the elevated level of miR-631in SCC131 cells.

**Figure 1.**
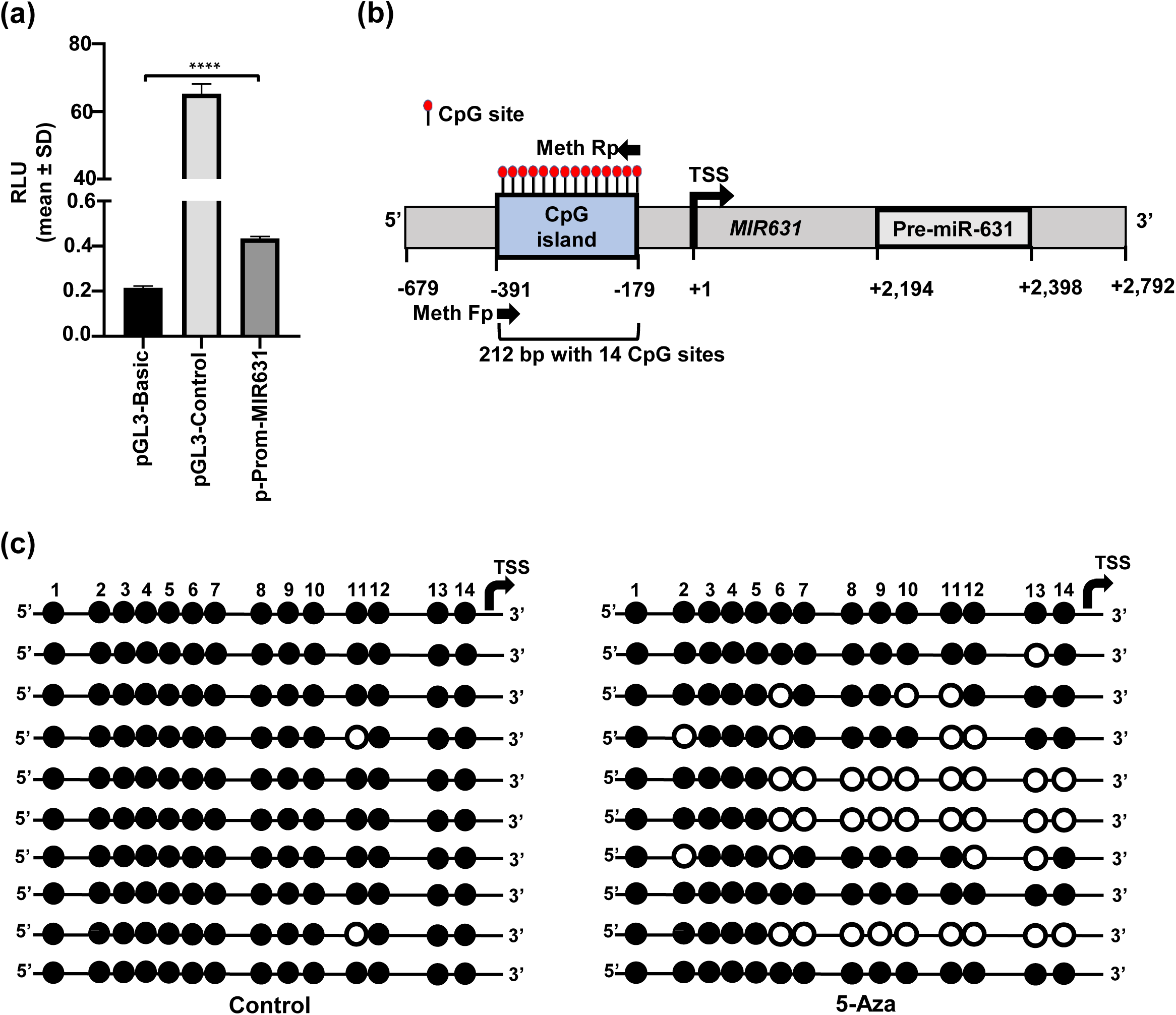
Characterization of the *MIR631* gene promoter and deciphering the mechanism of miR-631 upregulation in SCC131 cells. (a) The dual-luciferase reporter assay shows an increased luciferase activity for the putative promoter fragment cloned in pProm-MIR631. pGL3-basic is a promoter less vector, whereas pGL3-control harbors the SV40 promoter and was used a positive control. Each bar reflects the average result of three technical repeats performed in the dual-luciferase reporter assay (n=3). ****, *p*<0.0001. (b) Schematic representation of the *MIR631* gene promoter with a 212 bp long CpG island harboring 14 CpG sites. Pre-miR-631 represents the location of the precursor sequence of miR-631. (c) Schematic representation of the methylation pattern of the CpG island. Filled circles represent methylated CpG sites, whereas empty circles represent non-methylated CpG sites. Each line represents a single TA clone. *Abbreviations:* RLU, Relative light unit; Meth Fp, Methylation specific forward primer; and, Meth Rp, Methylation specific reverse primer.

### *In silico* identification and qRT-PCR analysis of putative gene targets of miR-631

miRs are known to exert their effects through target genes. To select a target gene for miR-631, five target gene prediction tools were used: TargetScan, miRwalk, DIANA, miRabel, and miRDB (**Table S1**). We focused on identifying target genes that were commonly predicted by these tools **(Table S1)** such as *AKR1B1* (Aldo-keto reductase family member B1), *OASL* (Oligo adenyl synthase), *RAP1B* (Rab associated protein 1B), *RAB11A* (Ras GTPase family member 11A) and *PTPRE* (Protein tyrosine phosphatase receptor). *PTPRE* is already a known target of miR-631. We also considered factors such as the number of target sites present in the 3’UTR of each target gene, predicted free energy of binding between miR-631 and the target gene, and accessibility of the target site, using bioinformatics tools like STARFold [https://sfold.wadsworth.org/cgi-bin/], and UNAFold [https://www.unafold.org/] (**Table S2).** Finally, we selected two potential target genes for further investigation: *RAB11A*, and *RAP1B*. To determine if miR-631 regulates these genes, we transfected SCC131 cells with pmiR-631 and the vector individually and determined their levels (**Fig. 2a, S5)**. The outcomes indicate that miR-631 regulates the transcript and protein levels of *RAB11A* (**Fig. 2a**). miR-631 does not regulate the transcript level of *RAP1B* (**Fig. S5**). As expected, *PTPRE* was found to be regulated by miR-631 **(Fig. S5)**. To investigate whether miR-631 exerts a dose-dependent effect on *RAB11A*, SCC131 cells were transfected with varying quantities of pmiR-631, and the levels of both RAB11A protein and transcript were evaluated (**Fig. 2b**). The data revealed a dose-dependent reduction of RAB11A levels (**Fig. 2b**). Since miR-631 was found to be high in 5-Azacytidine treated SCC131 cells, we examined the levels of RAB11A in this condition (**Fig. 2c**). As expected, RAB11A levels were downregulated in cells treated with 5-Azacytidine as opposed to cells treated with DMSO, suggesting that miR-631 regulates the levels of *RAB11A* (**Fig. 2c**).

**Figure 2.**
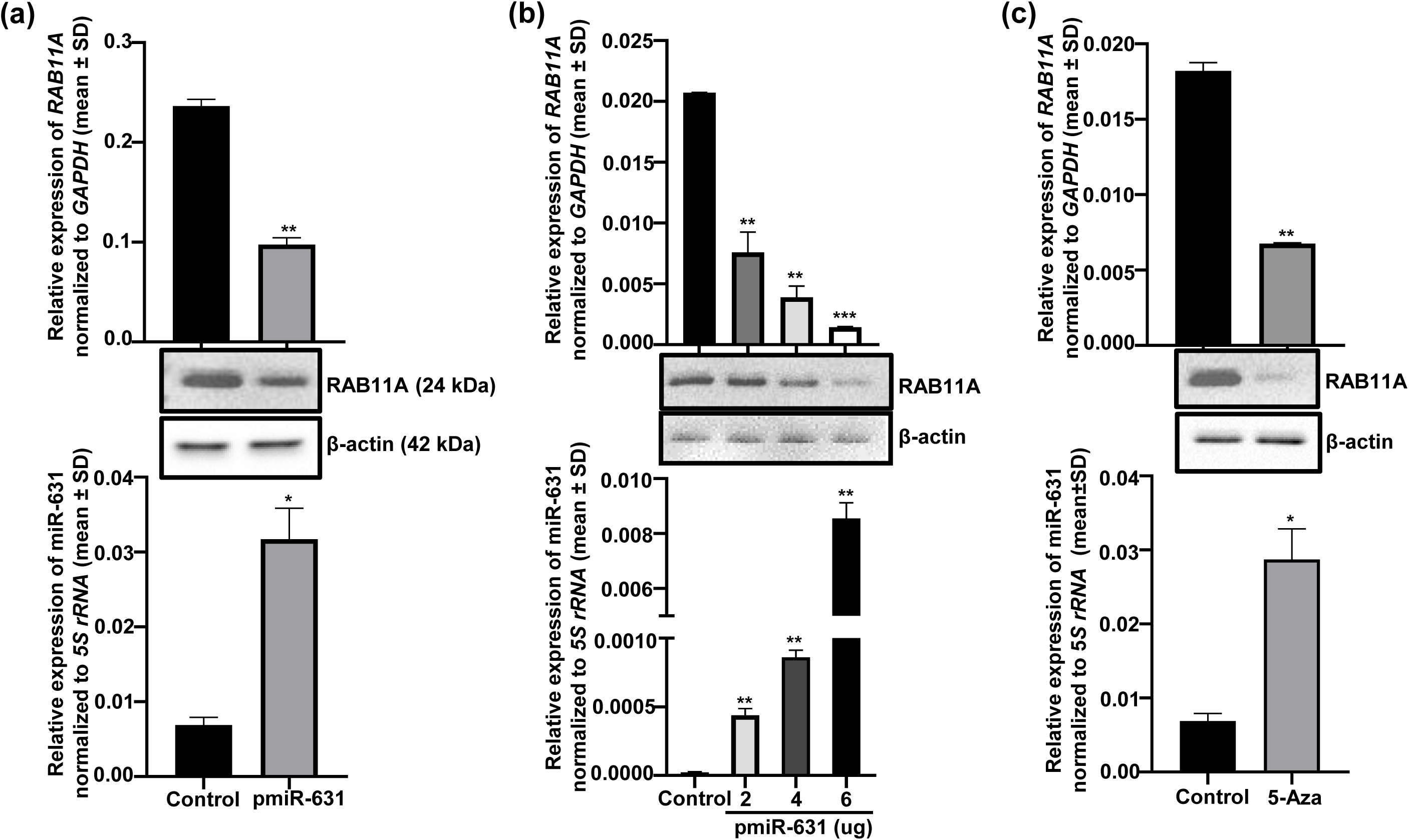
*RAB11A* is a potential target gene for miR-631. (a) Transfection of SCC131 cells with the pmiR-631 construct to induce miR-631 overexpression results in a decrease in protein and transcript levels of RAB11A (n=2). (b) The dose-dependent regulation of protein and transcript levels of RAB11A with increasing quantities of pmiR-631 in SCC131 cells (n=2). (c) Following 5-Azacytidine exposure of SCC131 cells, a decrease in both RAB11A transcript and protein levels was observed, accompanied by a concomitant increase in miR-631 expression (n=2). Both sets of samples in Fig. 2a, 2b, and 2c are processed in parallel and the full uncropped blot images are given in supplementary fig. S15. Each bar in the qRT-PCR data represents the mean of two technical replicates (n=2). *, *p*<0.05; **, *p*<0.01; ***, *p*<0.001.

### Confirmation of the direct interaction between miR-631 and *RAB11A*

A bioinformatics analysis predicted a target site (TS) within *RAB11A* 3’UTR for the seed region (SR) of miR-631 (**Fig. 3a**). Notably, the predicted TS showed conservation across different species (**Fig. 3a).** We employed a dual-luciferase reporter assay to confirm the direct binding of miR-631 to the *RAB11A* 3’UTR and therefore generated the following constructs in the pMIR-REPORT^TM^ vector: p3’UTR-RAB11A-S with the *RAB11A* 3’UTR in a sense-orientation, p3’UTR-RAB11A-M harboring the *RAB11A* 3’UTR with mutated TS, and p3’UTR-PTPRE-S with the *PTPRE* 3’UTR in sense-orientation. We also generated pmiR-631-M construct in pcDNA3-EGFP vector where the SR was mutated by the site-directed mutagenesis (**Fig. 3b**). We co-transfected different *RAB11A* 3’UTR constructs (e.g., p3’UTR-RAB11A-S and p3’UTR-RAB11A-M) separately with the vector, pmiR-631 or pmiR-631-M in SCC131 cells and determined the relative luciferase signal. Cells co-transfected with p3’UTR-RAB11A-S and pmiR-631 exhibited markedly lower luciferase activity in comparison to those with p3’UTR-RAB11A-S and the control plasmid pcDNA3-EGFP (**Fig. 3c**). Cells co-transfected with p3’UTR-RAB11A-M and pmiR-631 displayed luciferase activity similar to that observed in cells transfected with p3’UTR-RAB11A-S and pcDNA3-EGFP, attributed to the lack of a functional target site within the RAB11A 3’UTR in the mutant p3’UTR-RAB11A-M construct. (**Fig. 3c**). Likewise, cells co-transfected with p3’UTR-RAB11A-S and the mutant pmiR-631-M construct exhibited no change in luciferase activity relative to those with p3’UTR-RAB11A-S and pcDNA3-EGFP (**Fig. 3c**), owing to the absence of the seed region in the pmiR-631-M. As expected, the cells transfected with p3’UTR-PTPRE-S and pmiR-631 showed a notable reduction in luciferase signal when contrasted with those expressing p3’UTR-PTPRE-S and pcDNA3-EGFP (**Fig. 3c**). These findings indicate that the seed region (SR) of miR-631 binds directly to the TS in the *RAB11A* 3’UTR.

**Figure 3.**
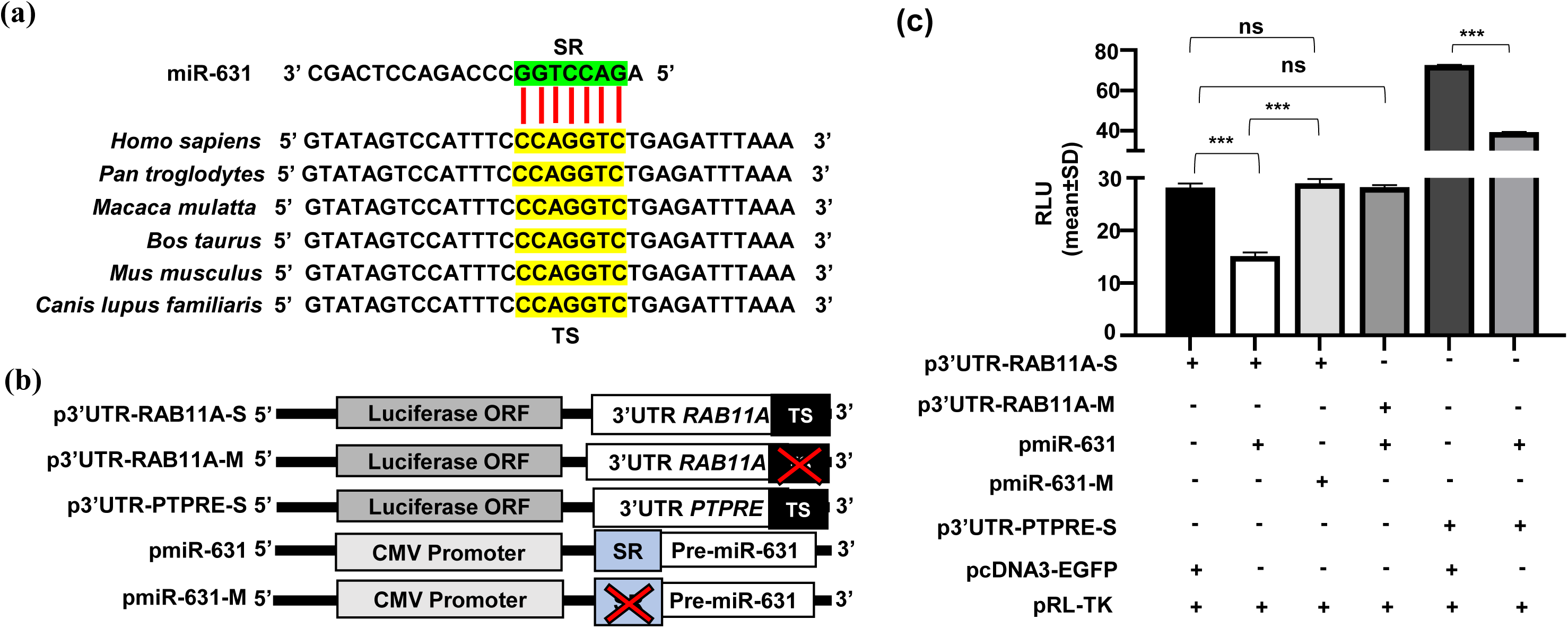
Confirmation of the interaction between miR-631 and *RAB11A*. (a) Presence of the putative TS in the 3’UTR of *RAB11A* conserved across different species for the SR of miR-631. (b) Schematic representation of the reporter gene constructs utilized in the luciferase assay. Abrogated TS or SR in a plasmid construct is indicated by X. (c) The dual-luciferase reporter assay in SCC131 cells co-transfected with different constructs. Note, a decrease in relative luciferase units was observed in cells co-transfected with p3’UTR-RAB11A-S and pmiR-631 compared to those with the vector plus pmiR-631. Each bar represents the mean of three technical replicates (n=3). ***, *p*<0.001. ns is when *p*>0.05. *Abbreviations:* TS, Target Site; and SR, Seed Region.

### Overexpression of *RAB11A* increases proliferation of OSCC cells

Since miR-631 reduces proliferation of both SCC131 and SCC084 cells, we wanted to determine the role of its target gene *RAB11A* on cell proliferation. To investigate this, the pRAB11A construct harboring the entire ORF of *RAB11A* and the cloning vector pcDNA3.1(+) were transiently transfected separately in SCC131 and SCC084 cells and the cell proliferation was determined, using the trypan blue dye exclusion method. The results revealed an enhanced proliferation of OSCC cells transfected with pRAB11A in comparison with those transfected with the vector **(Fig. S6a).** The same hypothesis was further tested using the BrdU incorporation assay, and the absorbance values indicated that both SCC131 and SCC084 cells transfected with pRAB11A showed increased cell proliferation in comparison with those transfected with the vector, confirming the pro-proliferative role of *RAB11A* **(Fig. S6b)**.

### Inverse correlation of miR-631 and *RAB11A* levels across various cancer cell lines and OSCC patient samples

To biologically interpret the relevance of miR-631/*RAB11A* interaction, next step was to evaluate the transcript levels of both miR-631 and *RAB11A* in various cancer cell lines using qRT-PCR. In general, an inverse correlation in the levels of miR-631 and its target gene *RAB11A* was observed among the cell lines **(Fig. S7)**.

We also checked the miR-631/*RAB11A* levels in 36 OSCC patient samples and their matched normal tissues by qRT-PCR **(Fig. S8)**. The miR-631 level was downregulated in 11/36 patient samples [viz., patient samples 50, 52, 55, 56, 62, 64, 68, 72, 82, 86, and 90], and its level was upregulated in 17/36 samples [viz., patient samples 38, 41, 54, 58, 61, 66, 69, 71, 76, 77, 78, 79, 80, 81, 87, 88, and 89]. The *RAB11A* level was upregulated in 13/36 patient samples [viz., 52, 55, 56, 57, 62, 64, 65, 68, 70, 76, 85, 89, and 90], whereas its levels was downregulated in 20/36 patient samples [viz., patient samples 38, 41, 50, 58, 61, 66, 69, 71, 72, 74, 75, 77, 78, 80, 81, 82, 83, 86, 87, and 88]. In 8/36 samples, miR-631 levels did not show any change [viz., patient samples 57, 63, 65, 70, 74, 75, 83, and 85]. The levels of *RAB11A* did not show any change in 3/36 patient samples [viz., patient samples 54, 63, and 79]. miR-631 was downregulated, whereas *RAB11A* was upregulated in 7/36 samples [viz., patient samples 52, 55, 56, 62, 64, 68, 90]. miR-631 showed upregulation and *RAB11A* showed downregulation in 14/36 samples [viz., patient samples 38, 41, 58, 61, 66, 69, 71, 74, 77, 78, 80, 81, 87, and 88]. Overall, 21/36 (58.33%) patient samples (viz. patient samples 38, 41, 52, 55, 56, 58, 61, 62, 64, 66, 68, 69, 71, 74, 77, 78, 80, 81, 87, 88 and 90) demonstrated an inverse relationship between the levels of miR-631 and *RAB11A* **(Fig. S8)**, indicating the biological significance of their interaction.

### Expression of *RAB11A* at the protein level is regulated by the presence of its 3’UTR

As demonstrated earlier (**Fig. 3**), the reduction in *RAB11A* levels results from the direct binding of miR-631 to *RAB11A* 3’UTR. To examine whether RAB11A protein levels are influenced by the inclusion or exclusion of its 3’UTR, we employed three constructs: (i) pRAB11A containing the entire ORF of *RAB11A* without the 3’UTR, (ii) pRAB11A-UTR-S which includes the complete ORF along with the 3’UTR in sense orientation, and (iii) pRAB11A-UTR-M with the full ORF and a mutated 3’UTR lacking a functional TS (**Fig. 4a**). These constructs with the vector [pcDNA3.1(+)] were transiently transfected separately in SCC131 and SCC084 cells, either alone or in combination with pmiR-631, and RAB11A protein levels were subsequently analyzed by Western blotting (**Fig. 4a, S9a)**. Cells co-transfected with pcDNA3.1(+) and pmiR-631 showed a decreased RAB11A protein level relative to those receiving only the vector, consistent with our earlier observation of miR-631 targeting the endogenous *RAB11A*. Co-transfection of pmiR-631 with pRAB11A in OSCC cells resulted in an elevated RAB11A levels compared to those transfected with the vector and pmiR-631, reflecting the absence of the 3’UTR in pRAB11A. However, co-transfection of pRAB11A-UTR-S with pmiR-631 led to a reduction in RAB11A levels when compared to the co-transfection of pRAB11A and pmiR-631, owing to the intact miR-631 binding site within the 3’UTR of pRAB11A-UTR-S. In contrast, higher RAB11A expression was observed in cells transfected with pRAB11A-UTR-M and pmiR-631 relative to those transfected with pRAB11A-UTR-S and pmiR-631, because of an abrogated TS in pRAB11A-UTR-M (**Fig. 4a, S9a).** Collectively, these findings indicate that miR-631 mediated regulation of RAB11A is modulated by the inclusion or exclusion of its 3’UTR.

**Figure 4.**
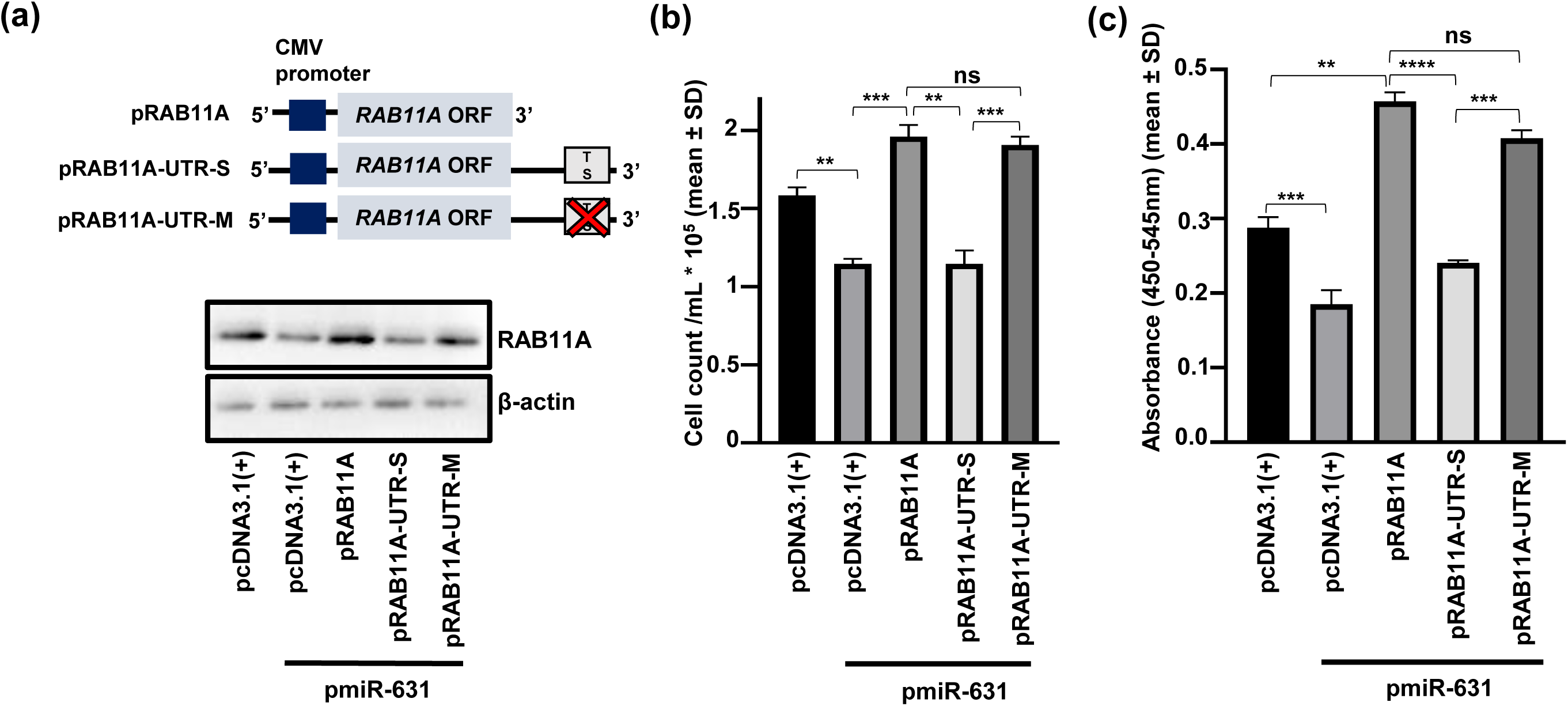
Regulation of *RAB11A* by miR-631 is mediated by its 3’UTR and the impact of this regulation on proliferation of SCC131 cells. (a) The RAB11A protein levels are dependent on the presence or absence of a functional target site in its 3’UTR for miR-631. Schematic diagrams of different *RAB11A* constructs are shown above the Western blots (n=2). Both sets of samples are processed in parallel and the full uncropped blot images are given in supplementary fig. S16. (b) The suppression of cell proliferation by miR-631, partly through targeting the 3’UTR of *RAB11A*, using the trypan blue dye exclusion assay. (c) The suppression of cell proliferation by miR-631, partly through targeting the 3’UTR of *RAB11A*, using the BrdU incorporation assay. Each datapoint is an average of three values (n=3). **, *p*<0.01; ***, *p*<0.001; ****, *p*<0.0001. ns is when *p*>0.05.

### Regulation of cancer hallmarks by miR-631, partly, via targeting the *RAB11A* 3’UTR

Next, we sought to explore the impact of miR-631 mediated regulation of RAB11A on cancer hallmarks, including cell proliferation, anchorage-independent growth, and apoptosis. We transfected pRAB11A, pRAB11A-UTR-S, and pRAB11A-UTR-M individually with pmiR-631 in SCC131 and SCC084 cells and checked cell proliferation using both trypan blue dye exclusion (**Fig. 4b, S9b**) and BrdU incorporation (**Fig. 4c, S9c**) assays. As anticipated, cells transfected with pmiR-631 and pcDNA3.1(+) exhibited a lower proliferation rate compared to those transfected with the vector alone (**Fig. 4b, 4c, S9b, S9c**), due to miR-631 targeting endogenous *RAB11A*. Additionally, a further reduction in cell proliferation was observed in cells transfected with pmiR-631 and pRAB11A-UTR-S in comparison with those transfected with pmiR-631 and pRAB11A, attributable to the presence of an intact target site in pRAB11A-UTR-S (**Fig. 4b, 4c, S9b, S9c**). In contrast, there was no significant change in proliferation between cells co-transfected with pmiR-631 and pRAB11A and those transfected with pmiR-631 and pRAB11A-UTR-M, consistent with the abrogated functional target site in pRAB11A-UTR-M. These findings collectively support the role of miR-631 in suppressing cell proliferation, in part, through direct interaction with the 3’UTR of *RAB11A*.

To examine the impact of miR-631-mediated regulation of RAB11A on the anchorage-independent growth of OSCC cells, we transfected SCC084 and SCC131 cells with the vector alone or co-transfected them with pRAB11A, pRAB11A-UTR-S, and pRAB11A-UTR-M with pmiR-631, followed by assessment of colony formation using the soft agar colony forming assay (**Fig. 5a, S10a)**. As anticipated, fewer colonies were formed in cells transfected with pmiR-631 plus vector in comparison with those transfected with the vector alone (**Fig. 5a, S10a)**. Cells transfected with pmiR-631 and pRAB11A-UTR-S showed a further decrease in colony formation relative to those transfected with pRAB11A and pmiR-631, due to binding of miR-631 to the TS in pRAB11A-UTR-S (**Fig. 5a, S10a)**. No significant difference in colony numbers was observed between cells transfected with pmiR-631 plus pRAB11A and those having pmiR-631 plus pRAB11A-UTR-M, since pRAB11A-UTR-M lacks a functional TS (**Fig. 5a, S10a)**. These findings indicate that miR-631 negatively regulates anchorage-independent growth, in part, by targeting the 3’UTR of *RAB11A*.To investigate whether miR-631 modulates apoptosis through targeting the 3’UTR of *RAB11A*, we transfected SCC084 and SCC131 cells with the vector alone or co-transfected them with pRAB11A, pRAB11A-UTR-S, and pRAB11A-UTR-M with pmiR-631, followed by the caspase-3 assay (**Fig. 5b, S10b)**. A significant increase in apoptotic cell death was observed in pmiR-631 plus vector co-transfected cells compared to those transfected with the vector alone (**Fig. 5b, S10b)**. The proportion of caspase-3 positive cells was markedly higher in the cells co-transfected with pmiR-631 plus vector relative to those co-transfected with pRAB11A plus pmiR-631 (**Fig. 5b, S10b)**. Furthermore, no notable difference in the proportion of caspase-3 positive cells was observed between cells co-transfected with pmiR-631 plus pRAB11A compared to those with pmiR-631 plus pRAB11A-UTR-M, as pRAB11A-UTR-M lacks a functional TS (**Fig. 5b, S10b)**. These results imply that miR-631 induces apoptosis in SCC131 and SCC084 cells, in part, through a direct interaction with the *RAB11A* 3’UTR.

**Figure 5.**
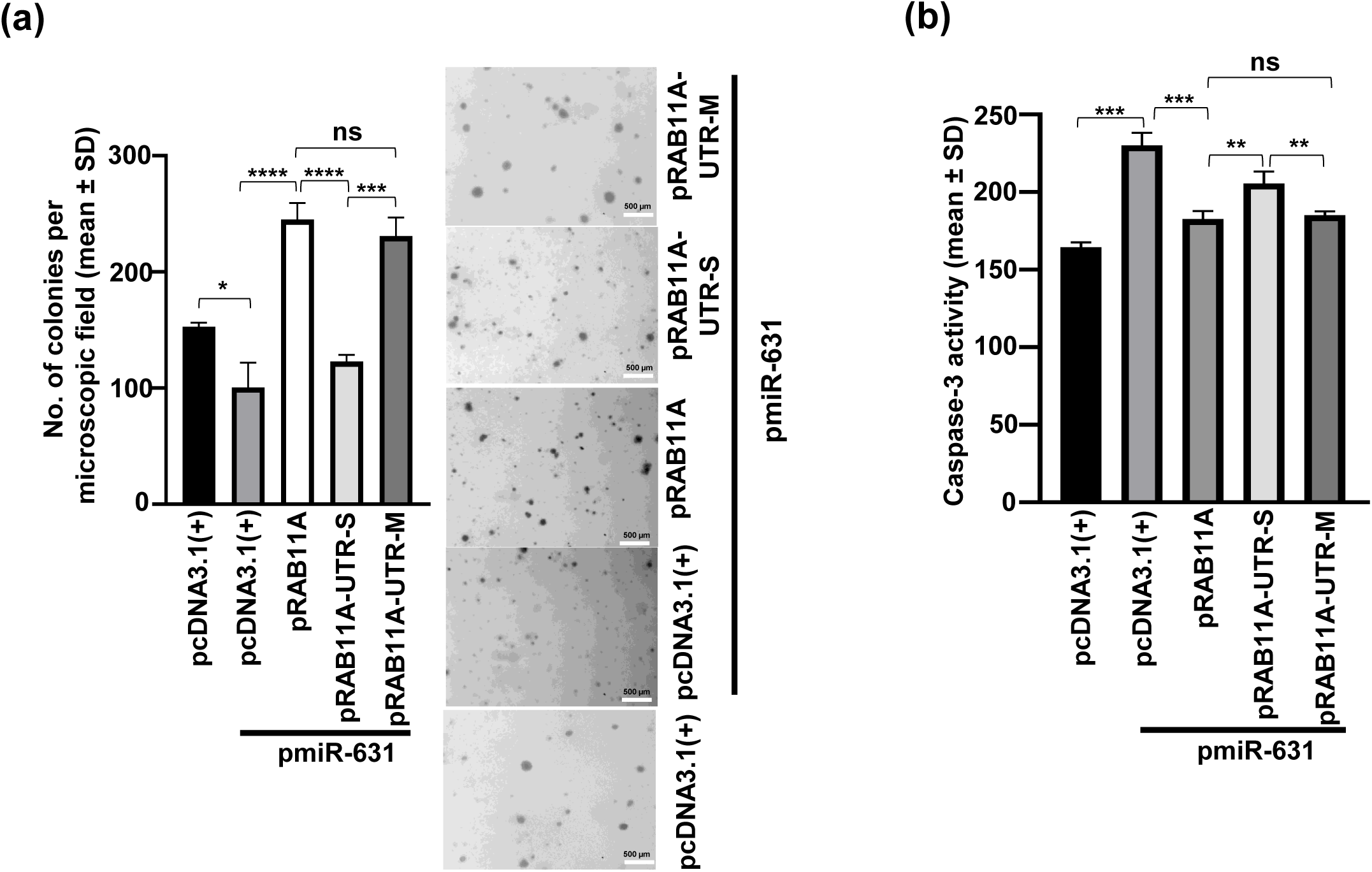
miR-631 modulates anchorage-independent growth and apoptosis, partly through targeting the 3’UTR of *RAB11A* in SCC131 cells. (a) The suppression of anchorage-independent growth by miR-631, partly through targeting the *RAB11A* 3’UTR. Representative micrographs of colonies formed by SCC131 cells subjected to co-transfection with pmiR-631 and various *RAB11A* constructs or the vector, obtained from the soft agar colony formation assay, are displayed on the right. (b) The positive regulation of apoptosis by miR-631, partly through targeting the *RAB11A* 3’UTR. Each datapoint is an average of three values (n=3). **, *p*<0.01; ***, *p*<0.001; ****, *p*<0.0001. ns is when *p*>0.05.

### Role of the miR-631/*RAB11A* axis in regulating Wnt signaling in OSCC cells

To gauge the activation of Wnt signaling, a gold-standard reporter assay employing the TOPflash reporter construct (TCF Reporter Plasmid) was used. This construct incorporates a minimal fos promoter coupled to TCF-binding sites upstream to the firefly luciferase gene. When the Wnt signaling is activated, the luciferase activity increases due to the greater amount of beta-catenin binding to the TCF site. In our study, we assessed the luciferase activity following the overexpression of pRAB11A or pmiR-631 in both SCC131 and SCC084 cells **(Fig. S11)**. The results revealed a significant increase in the luciferase activity with RAB11A overexpression **(Fig. S11a)**, whereas miR-631 overexpression led to a reduced luciferase activity **(Fig. S11b)**.

To determine whether RAB11A-mediated Wnt signaling activation in OSCC cells is regulated by miR-631, we employed the same *RAB11A* constructs (i.e., pRAB11A, pRAB11A-UTR-S and pRAB11A-UTR-M). SCC131 cells were co-transfected with the TOPflash reporter plus pmiR-631 or TOPflash reporter plus vector and determined the luciferase activity. The findings showed a decreased luciferase activity in cells co-transfected with TOPflash reporter plus pmiR-631 relative to those having TOPflash reporter plus vector **(Fig. S12)**. A significantly increased luciferase activity was observed in cells transfected with TOPflash, pmiR-631 and pRAB11A relative those having TOPflash plus pmiR-631, suggesting that RAB11A enhances the Wnt signaling **(Fig. S12)**. However, the luciferase activity decreased in SCC131 cells co-transfected with TOPflash, pRAB11A-UTR-S and pmiR-631 relative to those with TOPflash, pRAB11A-UTR-M and pmiR-631 **(Fig. S12).** This occurs due to an abrogated miR-631 TS in pRAB11A-UTR-M. The above observations suggested that RAB11A contributes to OSCC progression by enhancing Wnt signaling. Conversely, miR-631 acts as a negative regulator of Wnt signaling, in part, by targeting *RAB11A.* Similar observation was also made in SCC084 cells **(Fig. S12)**.

### A synthetic miR-631 mimic inhibits tumor growth *in vivo*

As miR-631 showed reduced cell proliferation in OSCC, we aimed to investigate the efficacy of a synthetic miR-631 mimic in reducing OSCC xenografts in nude mice. To achieve this, the optimal dosage of a synthetic miR-631 mimic in SCC131 cells was determined. We first introduced increasing quantities of the miR-631 mimic in SCC131 cells through transient transfection and determined the levels of miR-631 and RAB11A **(Fig. S13)**. Notably, all three quantities of the mimic (i.e., 200 nM, 400 nM and 600 nM) demonstrated a significant decrease in both RAB11A transcript and protein levels, accompanied by a corresponding increase in miR-631 transcript levels **(Fig. S13)**. We chose 400 nM of the synthetic miR-631 mimic as the dosage for the subsequent experiments in nude mice **(Fig. S13)**.

To determine if the synthetic miR-631 mimic reduces OSCC xenograft size in nude mice, SCC131 cells were transfected separately with both miR-631 mimic and a mimic control. Post 24 hr of transfection, we injected equal number of cells from each transfection in both flanks of three female nude mice in each group, i.e., control mimic and synthetic miR-631 (**Fig. 6a**). Tumor growth was monitored for 23 days post-injection. As expected, there was a significant decrease in weight and volume of OSCC xenografts among the group treated with the synthetic miR-631 mimic compared to the control group (**Fig. 6b**). We also excised xenografts from the mice on Day 23 and evaluated the levels of miR-631 and RAB11A to determine whether the synthetic miR-631 mimic could reduce RAB11A protein and transcript levels. The data demonstrated that both RAB11A protein and transcript levels were decreased in OSCC xenografts treated with the miR-631 mimic compared to those treated with the control mimic (**Fig. 6c**). These findings further support the tumor suppressor role of miR-631 in OSCC.

**Figure 6.**
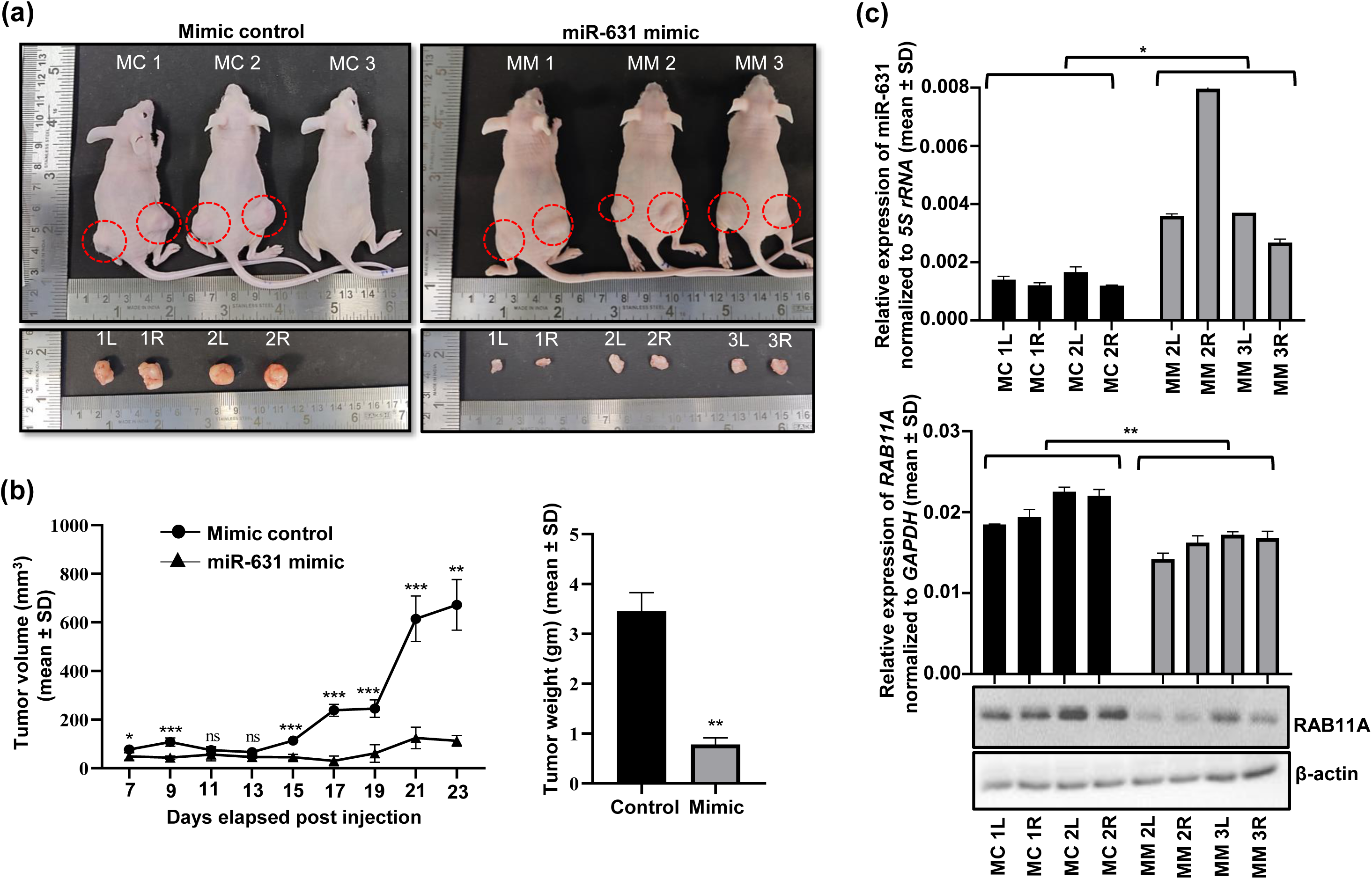
The impact of a synthetic miR-631 mimic on xenografts derived from SCC131 cells in nude mice. (a) Upper panel: Images of nude mice displaying tumor growth on Day 23 following injection of cells pre-treated with either a synthetic miR-631 Mimic (MM) or a Mimic Control (MC). Bottom panel: Excised xenografts from each mouse on Day 23. Xenografts are circled by red dash lines and labelled Left (L) and Right (R) according to the flanks they were excised from. Note, there was no tumor formation in nude mouse, MC3. (b) Impact of a synthetic miR-631 mimic on tumor volumes during a period of 7-23 days, and overall tumor weight of xenografts on Day 23. (c) qRT-PCR and Western blot analyzes of excised xenografts from the miR-631 mimic and the mimic control groups (n=4). Note, increased levels of miR-631 (Top panel) are correlated with reduced levels of RAB11A (Bottom panel) in xenografts with the miR-631 mimic as compared to those with the mimic control. Both sets of samples are processed in parallel and the full uncropped blot images are given in supplementary fig. S19. *, *p*<0.05; **, *p*<0.01; ***, *p*<0.001. ns is when *p*>0.05.

## Discussion

Studies have shown that miR-631 functions as a tumor suppressor in cancers [19–23]. For example, miR-631 targets *PTPRE* oncogene in hepatocellular carcinoma, whereas it targets the *ZAP70* oncogene in prostate cancer [19, 20]. In multiple myeloma, miR-631 downregulates *UBCH10* gene and it re-sensitizes doxorubicin-resistant chondrosarcoma cells by targeting the *APLN* gene [21, 22]. It also targets the *E2F2* gene in non-small cell lung carcinoma [23]. Our study has shown that miR-631 reduces proliferation of OSCC cells and reduces xenograft growth in nude mice, thus further strengthening its potential as a tumor suppressor (**Fig. 6, S2)**. Thus, miR-631 is an addition to a growing list of tumor suppressor microRNAs with roles in OSCC pathogenesis [10, 13, 14, 16].

The upregulation of miR-631 in SCC131 cells following the 5-Azacytidine treatment could be due to two possibilities: 1) demethylation of the *MIR631* gene promoter or 2) demethylation of the promoter of a transcriptional factor, which upon upregulation, activates the *MIR631* gene promoter. To investigate the first possibility, we characterized the *MIR631* gene promoter and assessed its methylation pattern in 5-Azacytidine-treated and untreated cells, using sodium bisulfite sequencing (**Fig. 1**). The analysis revealed decreased methylation at 14 CpG sites in the *MIR631* gene promoter in cells treated with 5-Azacytidine compared to untreated cells (**Fig. 1**), indicating that demethylation of the *MIR631* gene promoter causes its upregulation. Similarly, promoter demethylation of other tumor suppressor miRNAs, such as miR-617 and miR-198, has been shown to induce their upregulation in OSCC cells following 5-Azacytidine treatment [10, 14].

Each miR exerts its effect through target genes, but miR-631 has no known targets in OSCC. We shortlisted predicted target genes for miR-631 using five miR target gene prediction tools and focused on commonly identified ones across all databases. After a detailed bioinformatics analysis and *in vitro* assays, *RAB11A* was observed to be the direct target of miR-631 (**Fig. 3c**). We found that miR-631 reduces RAB11A levels in a dose-dependent manner (**Fig. 2b).** Since miR-631 shows upregulation post 5-Azacytidine treatment, we confirmed that the same treatment renders RAB11A levels down (**Fig. 2c**). *RAB11A* is a member of Ras associated GTPase family and is located on chromosome 15q22.31 **(Fig. S3)**. RAB11A upregulates YAP and Cyclins in NSCLC, whereas it upregulates p-AKT in gastric cancer [24, 25]. In the context of breast cancer, miR-320a, and miR-452 are shown to downregulate the expression of *RAB11A* [26, 27]. In ovarian cancer, RAB11A is downregulated by miR-193a-5p [28]. As an established oncogene having no characterized functions in OSCC, we looked further at the role of *RAB11A* in OSCC. Our results showed that RAB11A increases the rate of proliferation of OSCC cells, indicating its pro-proliferative nature **(Fig. S6)**.

To analyze the biological relevance of the interaction between miR-631 and *RAB11A*, we assessed their transcript levels in different cancer cell lines. A549 and U2Os with the lowest levels of miR-631 showed higher transcript levels of *RAB11A,* whereas Ln229 and MB-231 with higher levels of miR-631 showed reduced levels of *RAB11A* **(Fig. S7)**. Overall, an inverse correlation was detected in the levels of miR-631 and RAB11A across the cell lines **(Fig. S7)**. A similar analysis was also performed on 36 matched normal oral tissue and OSCC patient samples **(Fig. S8)**. Of the 36 patient samples, 21 (58.33%) showed an inverse correlation in the transcript levels of miR-631 and *RAB11A*, suggesting the biological relevance of their interaction **(Fig. S8)**. However, no inverse correlation in their transcript levels was observed in the rest of the OSCC samples **(Fig. S8)**. This could be due to tumor heterogeneity, redundancy of splicing factors, different etiopathogenesis, additional regulators of *RAB11A*, and the heterogeneous genetic composition of each patient [29, 30]. Inconsistencies in the levels of miRs and their target genes in patient samples were also reported earlier by Sarkar et al. [10], More et al. [13], Kaushik et al. [14], and Mallela et al. [31].

Given the potential for multiple miRNAs to regulate a single target, we investigated whether the observed regulation results from a direct interaction between miR-631 and *RAB11A*. It was found that miR-631 exerts its regulation on RAB11A levels only when the TS is present in the *RAB11A* 3’UTR (**Fig. 3, 4a, S9a**), and when the TS was mutated, RAB11A levels were not regulated by miR-631 (**Fig. 3, 4a, S9a**). Next, we wanted to check the effect of their interaction on a few cancer hallmarks such as cell proliferation, anchorage-independent growth, and apoptosis. Both trypan blue dye exclusion cell proliferation and BrdU incorporation assays showed that miR-631 regulates proliferation of both SCC131 and SCC084 cells, in part, via targeting the 3’UTR of *RAB11A* (**Fig. 4b, 4c, S9b, S9c**). Further, miR-631 reduced anchorage-independent growth of OSCC cells and RAB11A enhanced it (**Fig. 5a, S10a)**, in part, by targeting the 3’UTR of *RAB11A*. Additionally, miR-631 as a tumor suppressor increased cellular apoptosis in both SCC131 and SCC084 cells, in part, by targeting the 3’UTR of *RAB11A* (**Fig. 5b, S10b)**. Thus, our study supports the observations that RAB11A has potential in promoting cancer progression [24–28].

Next, we looked at the downstream effectors of miR-631/RAB11A axis. RAB11A promotes Wnt signaling and aids in cancer progression of pancreatic cancer [32]. As Wnt signaling has not been studied in OSCC so far, we checked the activation of Wnt signaling in OSCC cells, using the TOPFlash Wnt reporter plasmid [33]. The Wnt signaling was found to be upregulated in *RAB11A* overexpression **(Fig. S11)**. However, upon miR-631 overexpression, the Wnt signaling was significantly reduced, indicating that miR-631 downregulates Wnt signaling by binding to the 3’UTR of *RAB11A*, and this regulation was reduced when the TS of miR-631 was mutated **(Fig. S12)**. In OSCC, many Wnt ligands such as WNT1, WNT3A, WNT7B, WNT11, FZD7, TCF4, DKK1/2/3, and SFRP1/2/4/5 are known to be dysregulated [34]. However, the present study is the first one to explore the effect of RAB11A on Wnt signaling in the context of OSCC.

Since the discovery of microRNA molecules as potential targets for cancer therapy, studies have investigated their applications in cancer treatment [13, 14, 35]. Synthetic miR mimics are artificially designed small RNA molecules that replicate the function of endogenous microRNAs. In cancers, a tumor suppressor miR mimic can bind to its oncogene targets and suppress their action [36]. Since we showed miR-631 as a tumor suppressor in OSCC and it targets oncogene *RAB11A*, we wanted to assess its tumor suppressive potential in *in vivo* OSCC xenograft nude mouse model, using a synthetic miR-631 mimic. The results showed that the miR-631 mimic reduces tumor growth of OSCC xenografts in nude mice (**Fig. 6**). This suggested that miR-631 is indeed a tumor suppressor in OSCC, and it can be used for developing a potential therapeutic agent for OSCC. More et al. [13] have recently shown that the tumor suppressor miR-6741-3p mimic reduces size and weight of OSCC xenografts in nude mice. Sarkar et al. [10] and Kaushik et al. [14] demonstrated that synthetic mimics of miR-617 and miR-198 reduce the size and weight of OSCC xenografts in nude mice respectively.

In summary, our study is the first to show the tumor suppressive role of miR-631 in OSCC, demonstrate its ability to regulate key hallmarks of cancer and show suppression of tumor growth in nude mice, in part, by targeting the oncogene *RAB11A*. Our study further showed the role of miR-631 in modulation of the Wnt signaling. We have also identified promoter methylation as the primary cause of miR-631 downregulation in OSCC and observed an inverse correlation in the expression levels of miR-631 and *RAB11A* in various cancer cell lines and OSCC patient samples, suggesting the biological relevance of their interaction. Further, the present study highlights the potential of miR-631 as a future therapeutic target for OSCC.

## Materials and methods

### Cell lines

The UPCI: SCC131 and UPCI: SCC084 oral squamous cell carcinoma cell lines utilized in this study were generously provided by Dr. Susanne M. Gollin (University of Pittsburgh, Pittsburgh, PA, USA) [37], and were also acquired from the Leibniz Institute DSMZ (https://www.dsmz.de). A549 cells were received from the laboratory of Prof. Upendra Nongthomba, Department of Developmental Biology and Genetics, Indian Institute of Science (IISc), Bangalore. U2OS cells were supplied by Prof. Ganesh Nagaraju’s lab, Department of Biochemistry, IISc, Bangalore. MB-231 cells were obtained from Prof. Ramray Bhat’s lab, Department of Developmental Biology and Genetics, IISc, Bangalore. HeLa cells were sourced from Prof. Sachin Kotak’s lab, Department of Microbiology and Cell Biology, IISc, Bangalore. The LN229 cell line was provided by Prof. Kumaravel Somasundaram’s group, also from the Department of Microbiology and Cell Biology, IISc, Bangalore.

### *In silico* identification of targets for miR-631

Five gene target prediction programs, namely miRDB [38; https://mirdb.org/cgi-bin/search.cgi], DIANA-microT-CDS [39; https://dianalab.ece.uth.gr/html/dianauniverse/index.php?r=microT_CDS], miRwalk [40; http://mirwalk.umm.uni-heidelberg.de/], miRabel [41; http://bioinfo.univ-rouen.fr/mirabel/view/result.php?page=mir], and TargetScan [42, https://www.targetscan.org//] were used to identify target genes for miR-631 **(Table S1)**.

### Patient sample collection

This study used a total of 36 paired normal oral tissue and OSCC patient samples that were collected at the Kidwai Memorial Institute of Oncology (KMIO), Bangalore, from 13^th^ July, 2018 to 14^th^ November, 2018. The study was conducted with written informed consent from the patients and approval from the ethics committee of the KMIO (approval #KMIO/MEC/021/05.January.2018). All procedures complied with the ethical guidelines outlined in the Helsinki declaration. Tissue specimens, including resected oral tumor samples and matched adjacent normal tissues (collected from the distal surgical margin), were preserved in RNALater™ (Sigma-Aldrich, St. Louis, MO, USA) and stored at –80 °C until analysis. Tumor classification was performed using the TNM (Tumor, Node, Metastasis) staging system established by the International Union Against Cancer (UICC; https://www.uicc.org/). The summary of the patient cohort is tabulated in **Table S3**, and details of clinicopathological features for each patient sample is mentioned in **Table S4**.

### Total RNA extraction and qRT-PCR

Total RNA was extracted from samples (cells and tissues) using TRI Reagent™ (Sigma-Aldrich, St. Louis, MO, USA). RNA concentration was measured using a NanoDrop 1000 spectrophotometer (Thermo Fisher Scientific, Waltham, MA, USA). The Verso cDNA Synthesis Kit (Thermo Fisher Scientific, Waltham, MA, USA) was employed for synthesizing first strand cDNA from 1-2 µg of total RNA. The transcript levels of miR-631 were quantified according to the method described by Sharbati-Tehrani [43], using specific primers listed in S5 Table. *RAB11A* expression levels were assessed by qRT-PCR using transcript-specific primers **(Table S5)**.

### *In silico* identification of the putative *MIR631* gene promoter

The putative promoter sequence for the *MIR631* gene was retrieved from the DBTSS database [44; https://dbtss.hgc.jp/], and verified using promoter identifying algorithms such as Promoter 2.0 [45; https://services.healthtech.dtu.dk/services/Promoter-2.0/] and Alggen Promo [46; http://alggen.lsi.upc.es/].

### Plasmid constructs

To overexpress miR-631 in cells, the pmiR-631 construct was generated in the vector pcDNA3-EGFP, downstream to the CMV promoter. The promoter characterization of *MIR631* was conducted using constructs where putative promoter fragment retrieved from the DBTSS database was ligated upstream to the luciferase gene in the pGL3-Basic vector (Promega, Madison, WI, USA) to generate pProm-MIR631. Overexpression of *RAB11A* was carried out by cloning *RAB11A* ORF in the pcDNA3.1(+) vector to generate pRAB11A.

The direct binding of miR-631 to the *RAB11A* 3’UTR was confirmed by a dual-luciferase reporter assay. For this experiment, two constructs namely p3’UTR-RAB11A-S and p3’UTR-RAB11A-M were generated. The p3’UTR-RAB11A-S construct includes an intact 7-mer target site for miR-631 binding in the 3’UTR of *RAB11A* and was cloned in the pMIR-REPORT^TM^ vector (Invitrogen, Waltham, MA, USA). Whereas, the p3’UTR-RAB11A-M construct containing a mutated 3’UTR that lacks the miR-631 target site was generated by site-directed mutagenesis, using a standard laboratory protocol [47]. As a positive control, the 3’ UTR of the *PTPRE* gene was also cloned in the pmiR-REPORT^TM^ vector, and the construct was named as p3’UTR-PTPRE-S.

To further examine the ability of miR-631 to downregulate RAB11A protein levels via direct binding to the *RAB11A* 3’UTR, two additional constructs were generated: pRAB11A-UTR-S and pRAB11A-UTR-M. To this end, the *RAB11A* ORF that was cloned in the pcDNA3.1(+) vector (i.e., pRAB11A) was linked to either the *RAB11A* 3’UTR in a sense orientation (pRAB11A-UTR-S) or the mutated 3’UTR lacking the miR-631 binding site (pRAB11A-UTR-M). The constructs employed throughout the studies were confirmed through digestion by appropriate restriction enzymes, followed by Sanger sequencing on a 3730xl DNA Analyzer (Thermo Fisher Scientific, Waltham, MA, USA). Details of the primers used in generating the above constructs are given in **Table S6**.

### Cell culture, transient transfection, and reporter assays

Cells were cultured in Dulbecco’s Modified Eagle Medium (Sigma-Aldrich, St. Louis, MO, USA) supplemented with 10% (v/v) fetal bovine serum (Thermo Fisher Scientific, Waltham, MA, USA) and 1X antibiotic-antimycotic solution (Sigma-Aldrich, St. Louis, MO, USA) in a humidified incubator at 37°C with 5% CO_2_.

In each well of a 6-well plate, 2×10^6^ cells from either SCC131 or SCC084 cell line were seeded, and then transiently transfected with either a single construct or a combination of constructs, using the transfection reagent Lipofectamine™ 2000 (Thermo Fisher Scientific, Waltham, MA, USA) as per manufacturer’s instructions. After 48 hr of transfection, cells were processed to isolate total protein and RNA.

The Dual-luciferase reporter assay kit (Promega, Madison, WI, USA) was used for confirming the binding between the 3’UTR of a target gene and miR-631. It was also used for characterizing the gene promoter. For this purpose, required constructs were transfected in seeded cells (5×10^4^ cells/well). Upon 48 hr of transfection, cells were processed and readings were taken by the VICTOR X multilabel plate reader (PerkinElmer, Waltham, MA, USA). Control vectors like pRL-TK were used to normalize the transfection conditions [10, 12].

For the Wnt signaling analysis, the TOPflash reporter plasmid (Sigma-Aldrich, St. Louis, MO, USA) was transiently transfected along with other overexpression construct(s) in SCC131 cells (5×10^4^ cells/well) as described by Choi et al. [33]. For normalizing the transfection efficiency, control vectors such as pRL-TK were used (Sigma-Aldrich, St. Louis, MO, USA) [10, 12].

### 5-Azacytidine treatment of cells

SCC131 cells were grown for 24 hr and treated with 5-Azacytidine (Sigma-Aldrich, St. Louis, MO, USA) at a final concentration of 5 µM or the vehicle control DMSO (Sigma-Aldrich, St. Louis, MO, USA) separately for 5 days. Total RNA, protein, and genomic DNA were isolated from 5-Azacytidine- and DMSO-treated cells for further studies [10].

### Methylation analysis of the *MIR631* promoter

Genomic DNA isolated from 5-Azacytidine- and DMSO-treated cells were treated with 3.25 M sodium bisulphite (Sigma-Aldrich, St. Louis, MO, USA) to convert non-methylated cytosines to uracil, according to Frommer et al. [48]. The converted DNA was subsequently used in PCR, using the following primers specifically designed for bisulfite-treated DNA: forward primer (5’-AGATCGTTAAGTAGGTAAGGGGTG-3’) and reverse primer (5’-CACTCACAAAATAACCTCCTAAACC-3’). The 212 bp long amplified fragments were cloned in a TA-cloning vector pTZ57R (Fermentas, Waltham, MA, USA) and 10 clones from each treatment were sequenced on a 3730xl DNA Analyzer to identify methylated and unmethylated cytosines.

### Western blot hybridization

Lysates from cells were made using the CelLytic^TM^ M Cell Lysis Reagent (Sigma-Aldrich, St. Louis, MO, USA), while the CelLytic ^TM^ MT Mammalian Tissue Lysis Reagent (Sigma-Aldrich, St. Louis, MO, USA) was used to prepare lysates from oral tissue samples. After resolving the proteins in the lysates in a SDS-PAGE gel (Sodium dodecyl sulfate-Polyacrylamide gel electrophoresis), they were transferred to a PVDF membrane (Pall Corp., Port Washington, NY, USA), using a semi-dry transfer apparatus. The blotting membrane was then kept for blocking in 5% skimmed milk (Nestle, Gurgaon, India) in 1X PBST (Phosphate buffered saline tween 20).

After the blocking, three PBST washes were given, and the membrane was incubated with corresponding primary and secondary antibodies. An anti-RAB11A antibody (1:10,000 dilution, cat#ab128913) was purchased from Abcam (Cambridge, MA, USA). The anti-mouse β-actin (1:10,000 dilution, cat# A5441) purchased from Sigma-Aldrich (St. Louis, MO, USA) was used as a loading control. The anti-rabbit HRP-conjugated secondary antibody (1.5:5,000 dilution, cat# HP03) and anti-mouse HRP-conjugated secondary antibody (1:5,000 dilution, cat# HP06) were purchased from Bangalore Genei (Bangalore, India).

### Cell proliferation assays

The rate of proliferation of SCC131 and SCC084 cells transfected with an appropriate construct or co-transfected with a combination of constructs was assessed by the trypan blue dye exclusion assay as described by Karimi et al. [49]. Cell proliferation was also determined using a BrdU cell proliferation assay kit (Sigma-Aldrich, Germany), as described in Sarkar et al. [10].

### Apoptosis assay

Cellular apoptosis in cells transfected with the appropriate construct(s) was quantified using the CaspGLOW Fluorescein Active Caspase-3 Staining kit (BioVision, Milpitas, CA, USA), as per manufacturer’s protocol and the method outlined by Rather et al. [13].

### Soft agar colony-forming assay

Tumor cells can overcome anoikis to proliferate and form colonies in suspension within a semi-solid medium such as soft agar [50]. The anchorage-independent growth of cells co-transfected with a combination of constructs was analyzed by the soft agar colony-forming assay in 35 mm tissue culture dishes, following a standard laboratory protocol [10].

### *In vivo* assay for tumor growth

The effect of a synthetic miR-631 mimic on OSCC tumor growth was assayed in 6 weeks old female NU/J athymic nude mice. Briefly, 2×10^6^ SCC131 cells/well were seeded in two 6-well plates. Following overnight incubation of the cells in a CO_2_ incubator, each of the six wells of a plate was transfected with 400 nM of a synthetic miR-631 mimic. The same procedure was repeated for the mimic control. Post 24 hr of transfection, cells from each plate were harvested and pooled together. Further, 3×10^6^ transfected cells from each pooled group of cells were suspended in 200 µL of 1:1 dilution of DPBS and matrigel [Corning Matrigel Growth Factor Reduced Basement Membrane Matrix, LDEV-free, cat # 354230, NY, USA] in a 1.5 mL tube (6 tubes/pooled group). Cell suspension from each tube was then subcutaneously injected in each posterior flank of each of the three mice. Xenograft volumes were measured using a Vernier calliper.

Volumes of the OSCC xenografts were calculated using the following formula: V = L×W^2^×0.5, where L and W denote the length and width of the tumor, respectively [10, 13, 14]. Animals were photographed, and the OSCC xenografts were collected, photographed and weighed on Day 23. This study was carried out in accordance with the recommendations in the Guide for the Care and Use of Laboratory Animals of the National Institutes of Health and ARRIVE guidelines (https://arriveguidelines.org/). The animal experimental protocol was approved by the Indian Institute of Science’s Animal Ethics Committee (approval certificate# CAF/Ethics/936/2023). All nude mice were reared on a 12:12 hr light/dark cycle in proper sterilized cages with required food and water. All animal welfare protocols were strictly followed, including the use of anesthetics, special housing conditions, and efforts to minimize pain and distress. Xenograft growth and the health of animals were monitored daily by trained personnel to prevent undue suffering. The experimental period spanned from Day 7 to Day 23 post-injection. A humane endpoint was defined based on tumor size; once the largest xenograft exceeded the permitted size (on Day 23), the animals were euthanized the same day under sterile conditions using isoflurane (Sigma-Aldrich, St. Louis, MO, USA), followed by cervical dislocation, by trained staff to ensure minimal discomfort. No animals died naturally during the experiment. The miRIDIAN miRNA hsa-miR-631-Mimic (cat#C-300957-01-0020) and miRIDIAN miRNA Mimic Negative Control #1 (cat#CN-001000-01-20) were obtained from Dharmacon (Lafayette, CO, USA).

### Statistical analysis

A two-tailed student’s t-test was performed using the GraphPad PRISM8 software (GraphPad Software Inc., San Diego) to analyze the statistical significance of the difference between two data sets, with Welch’s correction wherever required. Differences with P-value 0.05 (*), P-value <0.01 (**), P-value <0.001 (***), and P-value <0.0001 (****) were considered statistically significant, whereas P-value >0.05 was considered as statistically non-significant (ns).

## Acknowledgements

We are grateful to the patients for providing the normal and tumor oral samples.

## Authorship contribution

AK. conceptualized the project and AK and HPM designed all the experiments. HPM. performed all the experiments. HPM. wrote the first draft of the manuscript. AK reviewed and edited the final draft of the manuscript. CG provided clinical samples for the study and RM helped with the sample collection. All authors reviewed the manuscript.

## Data availability statement

All data supporting the findings of this study are available within the manuscript or its supplementary files.

