## Supplementary Figures for "miR-631 suppresses oncogene *RAB11A* in oral squamous cell carcinoma"

*Joint corresponding authors:

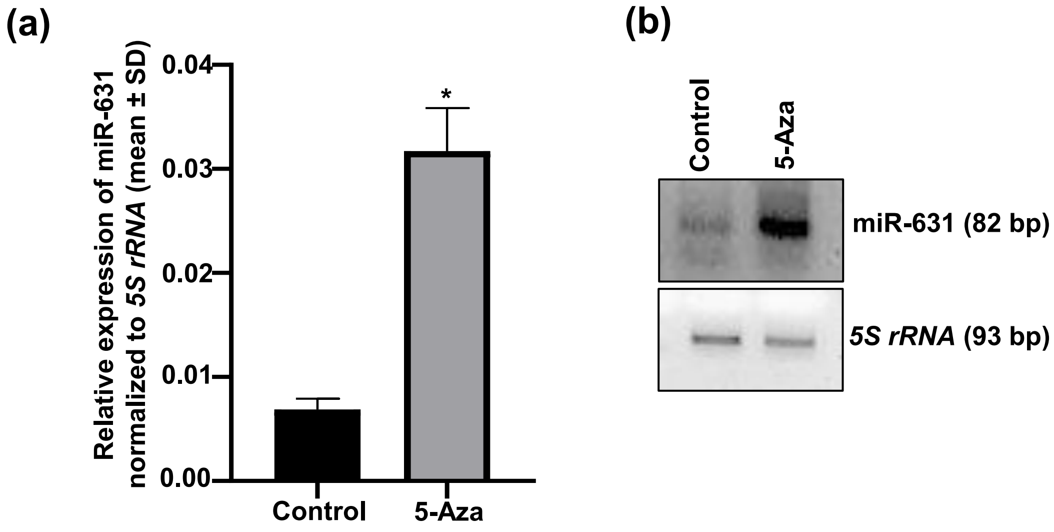


**Supplementary Figure S1.** Validation of miR-631 upregulation post 5-Azacytidine treatment. (a) The qRT-PCR analysis showing upregulation of miR-631 post 5-Azacytidine treatment of SCC131 cells. Each data point for qRT-PCR is an average of three technical replicates (n=3). (b) The semi-quantitative RT-PCR analysis showing upregulation of miR-631 in SCC131 cells following 5-Azacytidine treatment of SCC131 cells. *5S rRNA* was used as an equalizing control. *,$p$<0.05. *Abbreviation:* 5-Aza, 5-Azacytidine.

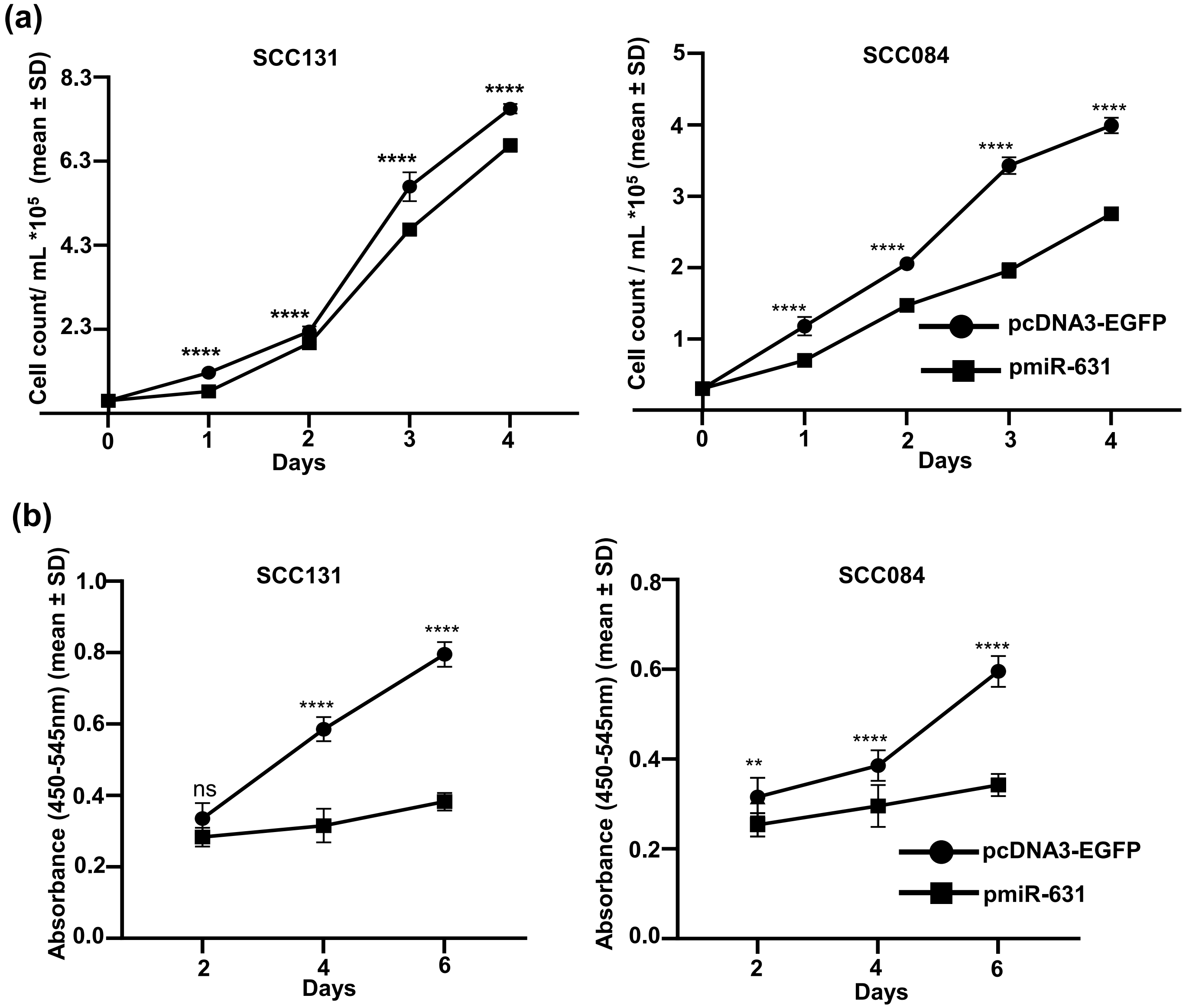


**Supplementary Figure S2**. Effect of miR-631 on OSCC cell proliferation. (a) The trypan blue dye exclusion assay to show anti-proliferative nature of miR-631 in SCC131 and SCC084 cells. Each data point of cell count represents an average of 6 values (n=3). (b) The BrdU incorporation assay to show the anti-proliferative nature of miR-631 in SCC131 and SCC084 cells. Each data point in BrdU absorbance values represents an average of three wells (n=3). **, $p<0.01$; ****, $p<0.0001$. ns is when $p>0.05$.

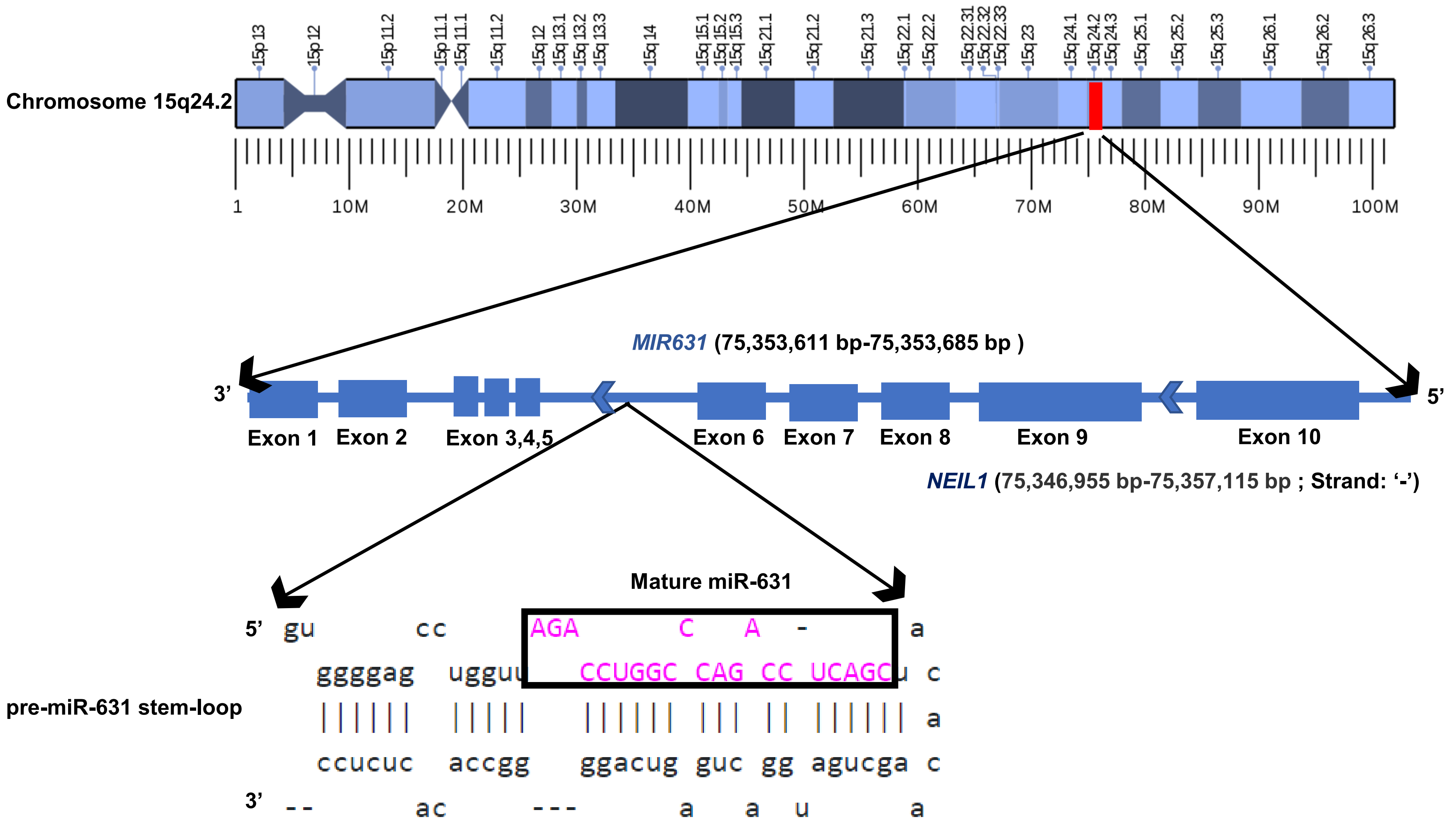


**Supplementary Figure S3.** Diagrammatic representation of the genomic location of miR-631 and pre-miR-631 stem-loop. The mature miR-631 sequences are highlighted in a black rectangle in the pre-miR-631 stem-loop.

TFIIB
**5'cacgagttcacctatgcactgatgc**cgcacaagggtgagtgctgggga**cg**gtggggccttcctgcagccctagacacctctctgcccttccctgagcccaaaccccagcaacctggcttcctgccccttctagggacaggggga
gggtagcccccattctccaccagcattgtttcctggtcgtaccccaggctctttccaggatgctggcgttatccaagctgcctacagcctaaacttcccccctgttggctctgccagcccccagcccagcgcccgccacctcctggagtgc
gttttccgtgtcttcacccgcggtcgtattggagac**cg**tcaagcaggcaaggggtggga**cg**tgctgggga**cg**ggttc**cg**aactgggcctgccct**cg**gcccctcccaccaga**cg**tgttcctctgcagg**cg**gagagcagccccc
ag**cg**c**cg**ct**cg**ctggtcctgaggctgtatgaggccca**cg**gcagcca**cg**tggactgctggctgcacttgt**cg**ctgc**cg**gttcaggaggccatcctgtgagtggcggtgtgggattggggcgggggtcgagtctcgccctcgcag
TFIIB TFII TSS
agccttcatggcccctcctgttctccagctgcgatctcttggagcgaccagaccctgctggccacttgacccttcgggacaaccgcctgaagctcacctttctcccttccaagtgctgtccctgttgctcgtgcttcagcctccgccacac
TFIIB TFIIB
tgagtccctggggctggggttttgtttgtagaaggctctggggactcctaatttctgcttccccagcctaaagcagggatcagtcttttcttgtggaataaatccttggatcgggaaccctgctgctcagtccccttagggcccatcctcacg
TFII
cttggggtgggggtgggaagtctggggtgggatggcttcccaggtatttggtgggtaggggcaagatcctaaggtacctgtgcgtgcagggtctgggttgcagcccccacaggagagactggggacctcgagccaccaaaagcca
gtgaagagaccatgagttgagaagcccctcccctgtgccagcaggtaaagggcctgggcctggggctggggagggggccgaagggtgggttttgcctggctcctgccaggatgagggggcagccctggaccctggagaatctgg
actggaggagctctggctcctgggctgcctggggccagggtctttgtaccacctctgaggggctgtccttgcagaggcaaactgtcctctgatagagtcagagccttgaggactgggatgggggctcagtttgccttgtgaggagctg
ggccttctgtcaccaggaccggacagcaggcccttcctacctggccatctcagaccaagaggtgccaagaagttcaaatgtatcagtttattaaaaatgcagcatttttcacatgagctttaaagatgtggaagatggggtacaattaaaa
ccatgagagttgtgcagggaacagccgtagggcctgtttgcaccttcagatattgcctgctcccaaaaattcagacccccagatgcagggcaagacaataagaaagggtgagtgcaagcaaggagagcctcctgctaagaggctga
ggtcccctctggttccaaggatgggatgtcagccttgaccttccggggtctgcagtggcctgggggaagaacaggcaagatggtcagctaagactccaggggctgccatggcagcctttctcccacctttgtacacgcgctcctaggc
caccatcccatcctctcctgtagagccagtttctgggagtgatgaggaaagcccaccagaggctgcctgtcgccccttcctcctccccttcttgggcactgtgggagcttctgggtcctgctggaggctggtcccctcaggccgctgggt
tgcagtcctcttaggaaggtctctctttgcccttcgtgtcctggaaggggccttgcttggaggcaaagcgtcctggatggaggagacacgggtccctgaggggctcagaaagcagttcagaggttggttagggcccaccactcccccc
gccctcttagggtaaggcagctgcctgggagggggcagccagggctgcagagtgcaggggaggaggcgggagcaggcagccatacctccactctgtcctcaggactcagctgtgtggccttggatttctttttgcgggacttgcgcc
ctgcaggacactggtgttggagttggagggtcctatcctgcccaggggtgactcccagggttgcagggggatagggtggagaagggtgctgtagcccttgcaggcgtgaagtcctttctgctctcttagcctattacattaggagtagct
tacctttgggtgccaacggtccaggatccccctaaaatgggatggggataattcaggaatcagcctgggttggcacaggggcggtattccttggagaggcaggactcacacacacccatccagatcagtgtagcttctcccttaggaa
gcctctaggacatcccccatgttagagtccacatcagcaaagctgctctgcccttggctactttcacttgggctacctgccttgggctacttccactagctgcaaccctgggacgcatgggagggggagggtgtgaccctcaggaacag
tgtggtccttggagggtctagacagaccctgagcatcaccaccccagttattgtgaccccacgtttccacccatcagcctcctggggtctctgcctgtgtgaacagtagggcccaacctggaaccagatggtacggccatgccggtcct
gcagggagctcatgcctggcatgccatagcagcgcagccaggctcgaaaggcagcaaagtcctcctccc**cgctctctgacccgtagcctttg**ccccctgtgggacagaggaacaggcagagatcagagggcaggctcaggttg
ggaggagtggggagcctggtt**agacctggcccagacctcagc**tacacaa**g**ctgatggactgagtcaggggccacactctccctcctctggtgatgtgacctcagctg**gtttcttcccactcggccatggg**tttcccatcctgga
gtgggattaagaatccttgtcctggccctgtgcagtggccacacctgtaatcccgacactttgggaagcttagatgggagaatccctaggggccaggagttcaataccagcctgggcaacatagggagaccctgtctctacaaaaaaa
atttttaacaattagcccggtgtggtggtgtgagcctgtagtcctagctacttagaggcagatgtgggaggatcacttgagcccagtttgaggctagtccaggcaacatggcaagatcctatctctaccaaaaatatatataaacgtgcgt
TFIID
gtgtgtgtgtgtgtgtgtgtgtataaatatataaaagaatccttgtcctgcct**gtctcatgggcagcttggaggaaac 3'**
TFIID

**Supplementary Figure S4.** Diagrammatic representation of the *MIR631* gene promoter sequence. Transcription start site (TSS) is marked with a curved blue arrow. The highlighted stretch of sequences in yellow color represents the CpG island analyzed for methylation pattern. The CpG sites are shown in bold letters. The binding sites for general transcription factors are highlighted in green. Precursor sequences for miR-631 are highlighted in grey color, whereas the red bold letters represent mature miR-631 sequence and black bold letters represent the primers.

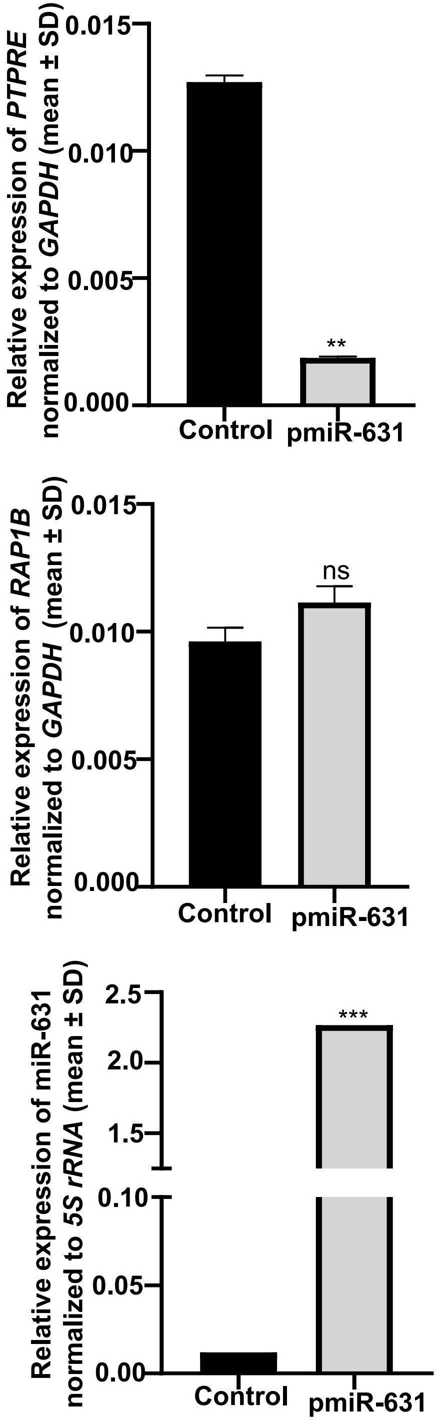


**Supplementary Figure S5.** Regulation of predicted target genes by miR-631. Overexpression of miR-631 in SCC131 cells causes downregulation of the target gene *PTPRE* and no change in the levels of another predicted target gene *RAP1B*. Each bar is an average of three technical replicates (n=3). **, $p<0.01$; ***, $p<0.001$. ns is when $p>0.05$.

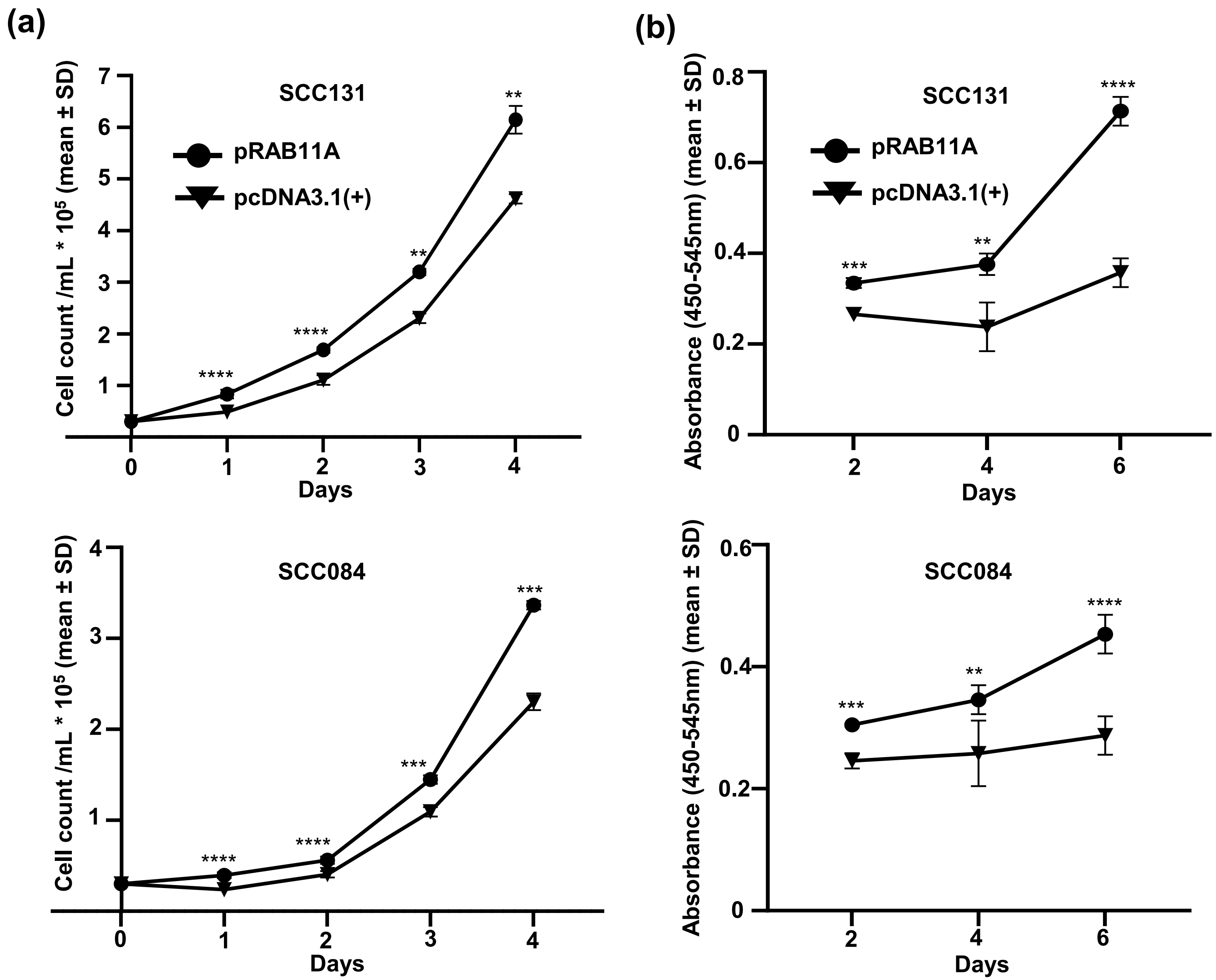


**Supplementary Figure S6.** The effect of *RAB11A* on cell proliferation. (a) Overexpression of *RAB11A* in SCC131 and SCC084 cells causes increased cell proliferation as shown by the trypan blue dye exclusion method. Each data point for cell proliferation is an average of six values (n=3). (b) Overexpression of *RAB11A* in SCC084 and SCC131 cells causes increased cell proliferation as shown by the BrdU incorporation method. Each data point of absorbance value is an average of three values (n=3).**, $p$<0.01; ***, $p$<0.001;****, $p$<0.0001.

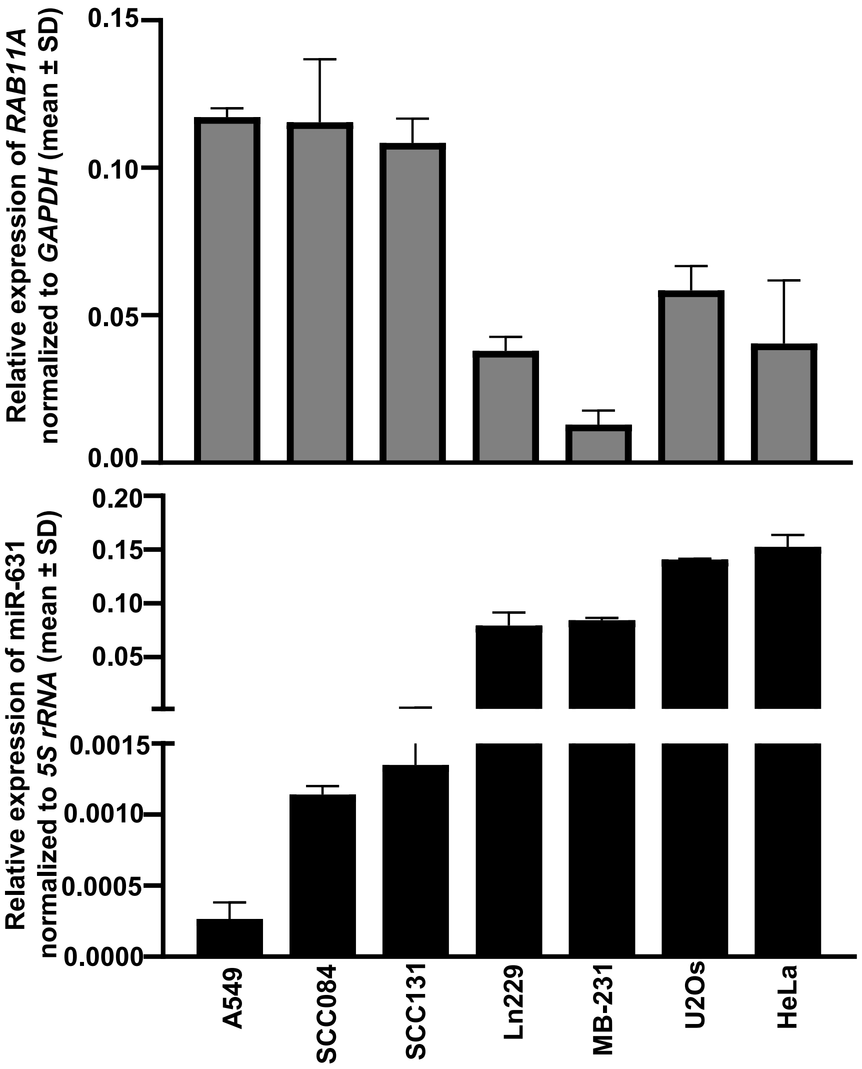


**Supplementary Figure S7.** Transcript levels of miR-631 and *RAB11A* in different cancer cell lines. In general, there appears to be an inverse correlation in the transcript levels of miR-631 and *RAB11A* across different cancer cell lines. Each bar is an average of two biological replicates (n=2).

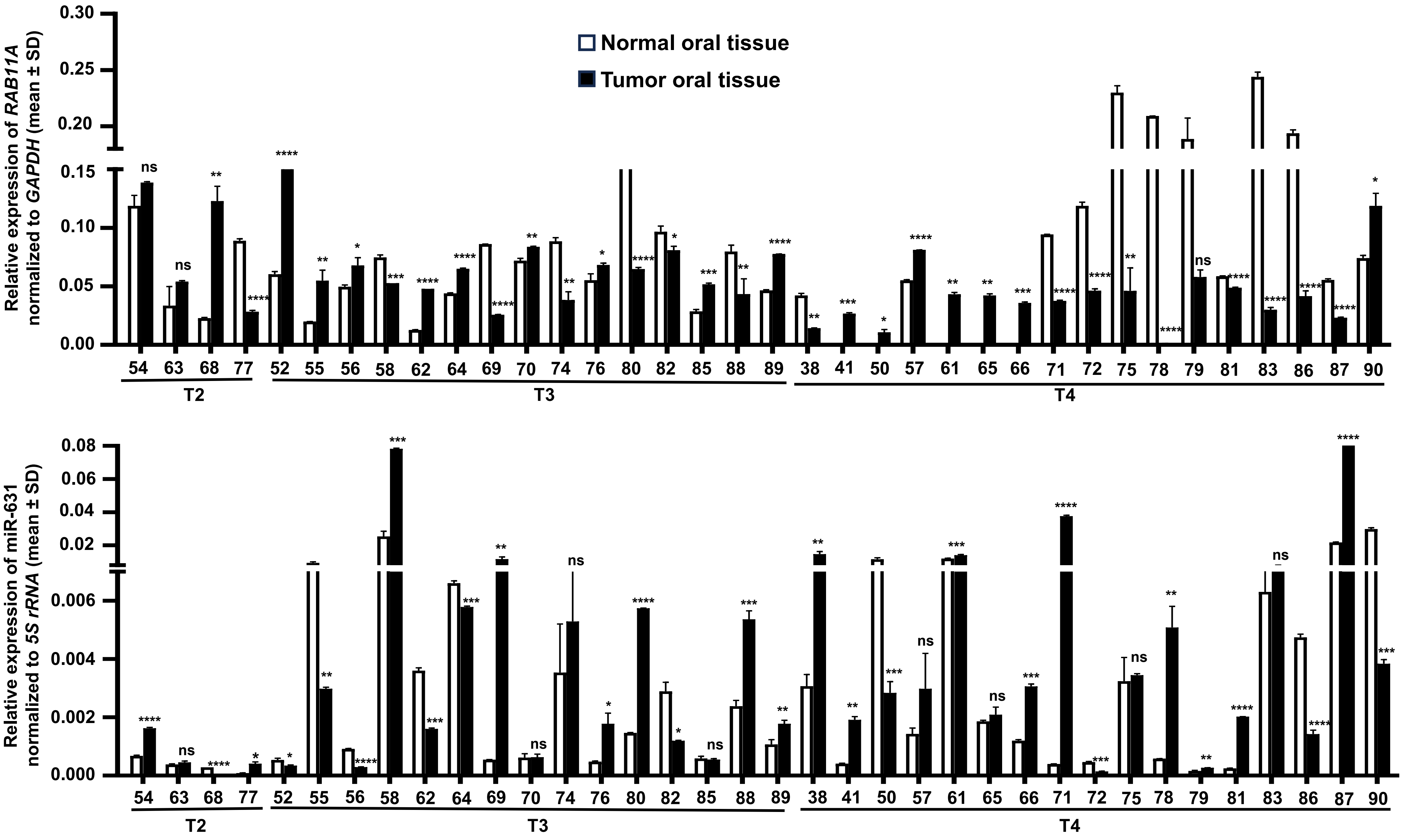


**Supplementary Figure S8.** Transcript levels of miR-631 and *RAB11A* in 36 matched normal oral tissue and OSCC patient samples. The T2, T3 and T4 represent the different stages of tumors, and the numbers along X-axis denote different patient numbers. Each qRT-PCR data is an average of 2 technical replicates (n=2). *, $p<0.05$; **, $p<0.01$; ***, $p<0.001$; ****, $p<0.0001$. ns is when $p>0.05$.

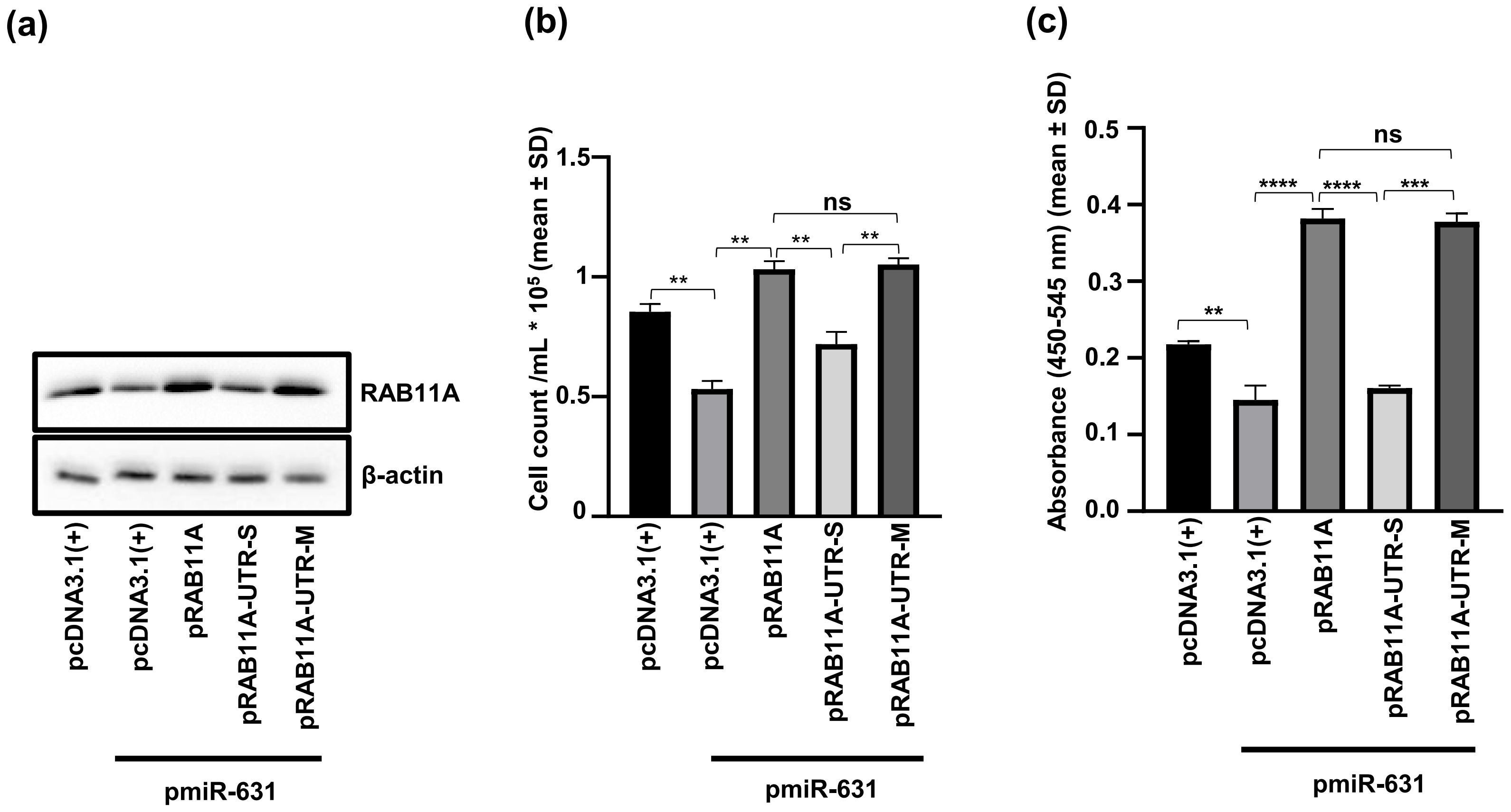


**Supplementary Figure S9.** The negative regulation of RAB11A by miR-631 is mediated by its 3'UTR and the impact of this regulation on proliferation of SCC084 cells. (a) The Western blot analysis shows that the RAB11A protein levels are dependent on the presence or absence of a functional TS in its 3'UTR for miR-631. Both sets of samples are processed in parallel and the full uncropped blot images are given in supplementary fig. S17. (b) The negative regulation of cell proliferation by miR-631, in part, via targeting the 3'UTR of *RAB11A*, using the trypan blue dye exclusion assay. (c) The negative regulation of cell proliferation by miR-631, in part, via targeting the 3'UTR of *RAB11A*, using the BrdU incorporation assay. Each datapoint is an average of three values (n=3). **, $p<0.01$, ***, $p<0.001$, ****, $p<0.0001$. ns is $p>0.05$.

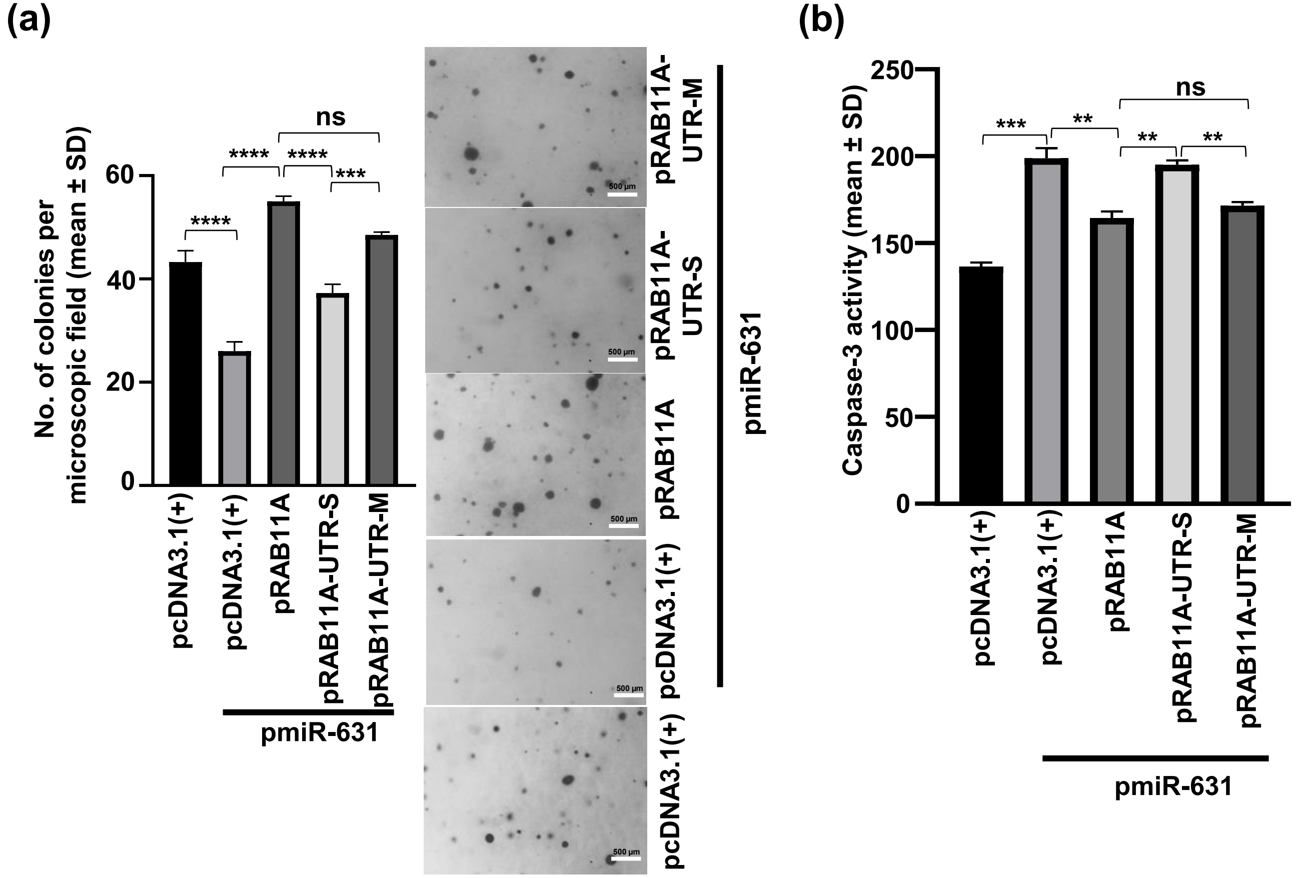


**Supplementary Figure S10.** Regulation of anchorage-independent growth and apoptosis by miR-631, in part, via targeting the 3'UTR of *RAB11A* in SCC084 cells. (a) The negative regulation of anchorage-independent growth by miR-631, in part, via targeting the 3'UTR of *RAB11A*. Representative microphotographs of the colonies for SCC084 cells co-transfected with pmiR-631 and different *RAB11A* overexpression constructs or vector control from the soft agar colony forming assay are shown on the right. (b) The positive regulation of apoptosis by miR-631, in part, via targeting the 3'UTR of *RAB11A*. Each datapoint is an average of three values (n=3). **, $p<0.01$, ***, $p<0.001$, ****, $p<0.0001$, ns is when $p>0.05$.

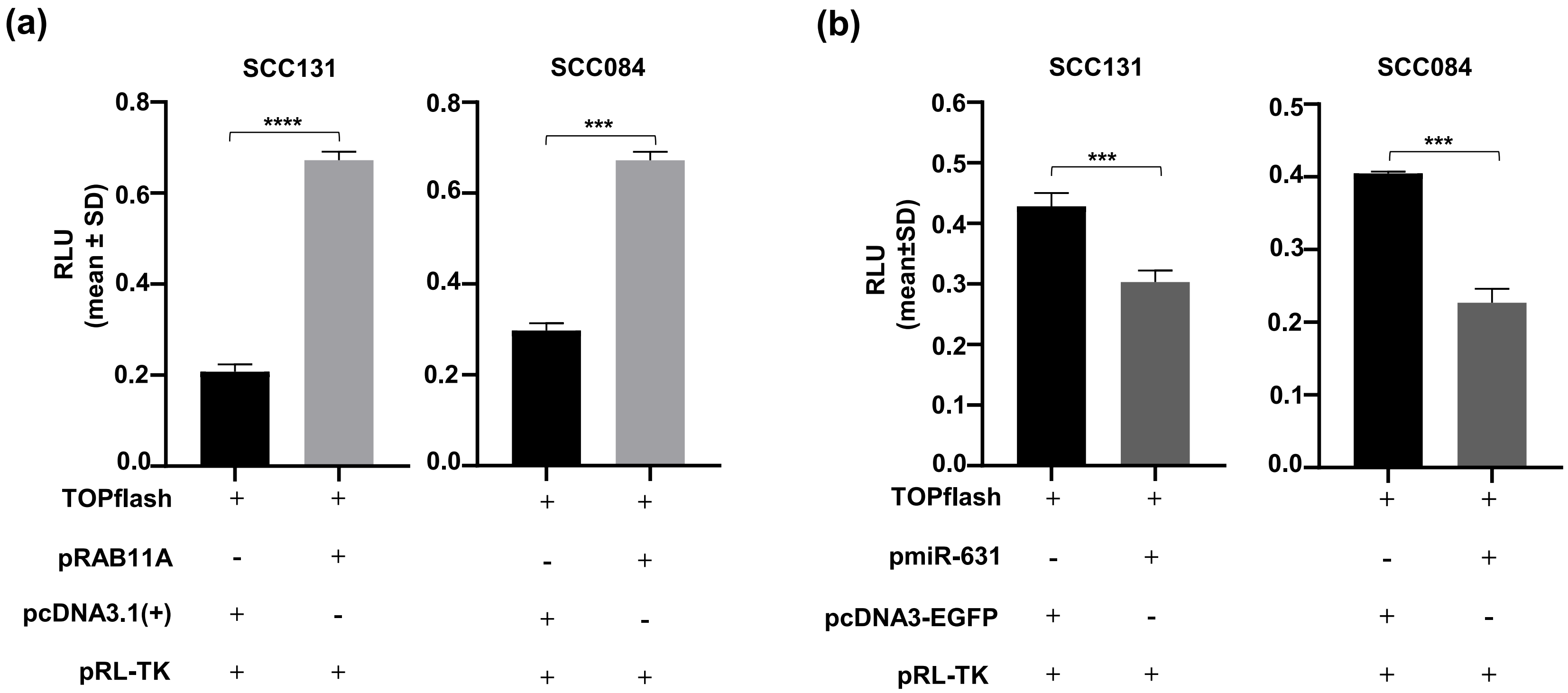


**Supplementary Figure S11.** RAB11A enhances Wnt signaling, while miR-631 negatively regulates it in SCC131 and SCC084 cells. (a) The TOPflash reporter assay to show the positive regulation of Wnt signaling following overexpression of RAB11A. (b) The TOPflash reporter assay to show the negative regulation of the Wnt signaling following overexpression of miR-631. Each bar is an average of three values (n=3). ***, $p<0.001$; ****, $p<0.0001$. *Abbreviation*: RLU, Relative light unit.

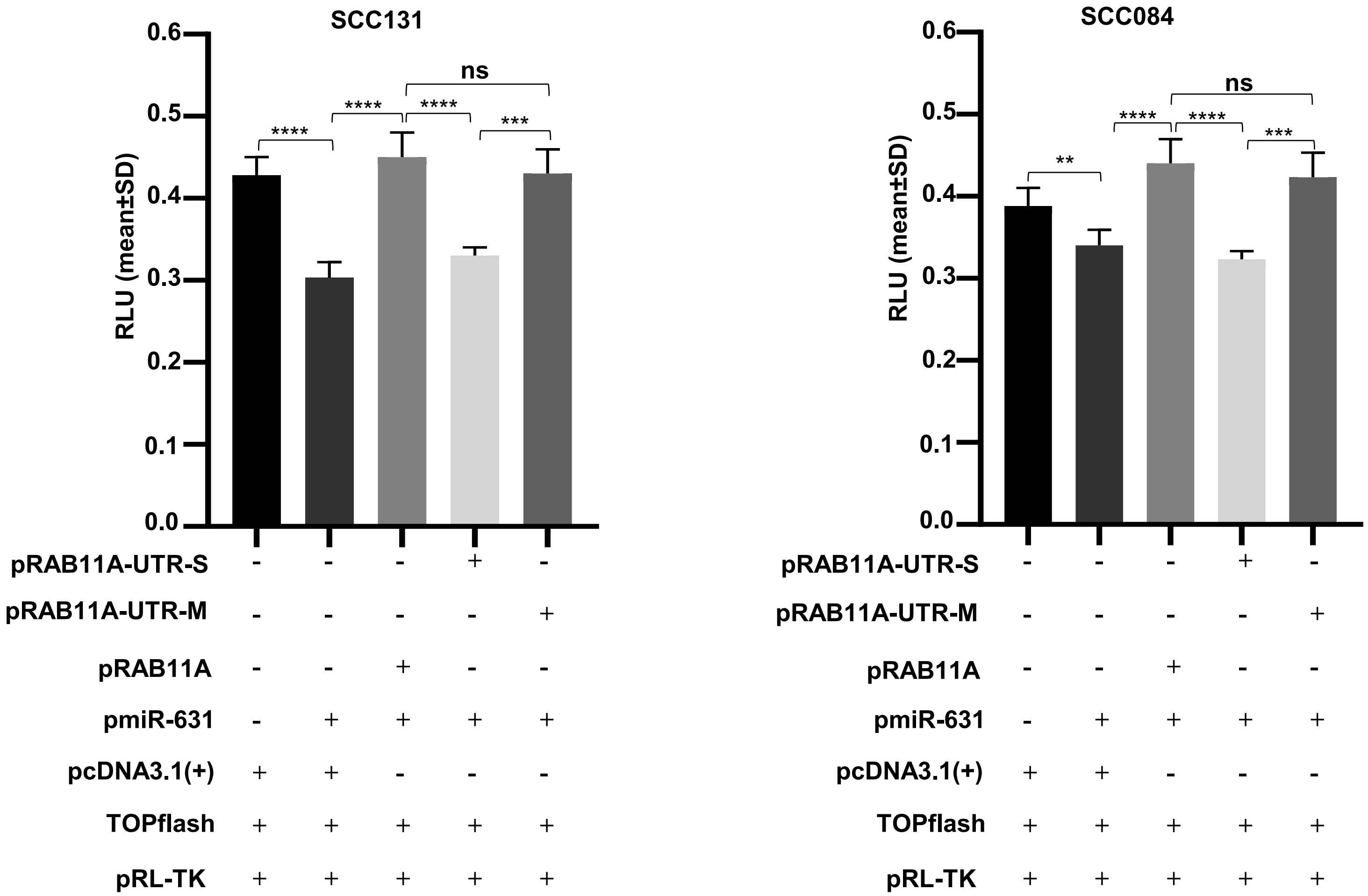


**Supplementary Figure S12.** miR-631 negatively regulates Wnt signaling, in part, by targeting the 3'UTR of *RAB11A* in SCC131 and SCC084 cells. The upregulation of Wnt signaling following overexpression of *RAB11A* is dependent on the presence or absence of its 3'UTR. Each bar is an average of three values (n=3). **, $p$<0.01; ***, $p$<0.001;****, $p$<0.0001. ns is when $p$>0.05.

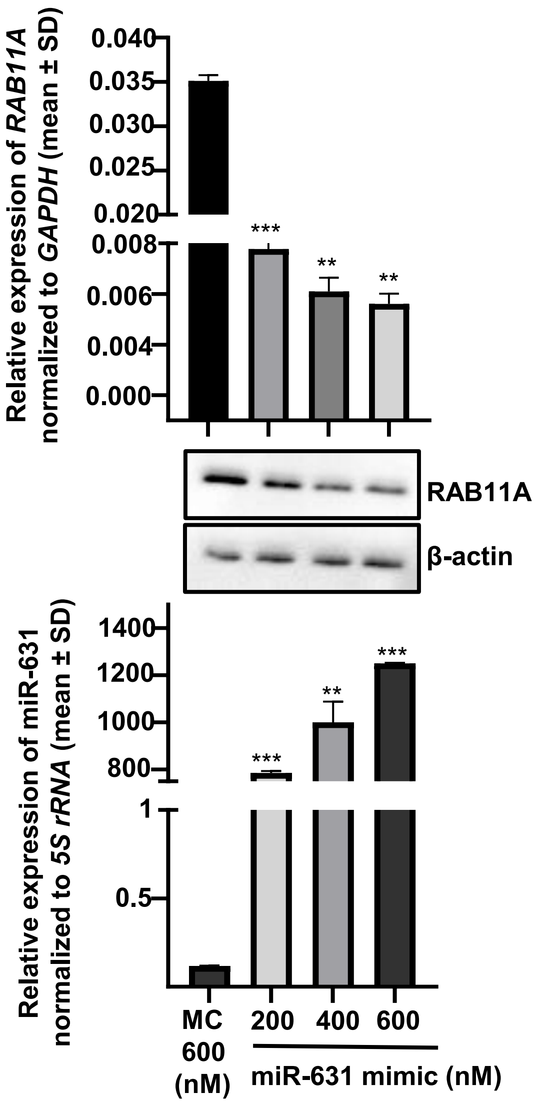


**Supplementary Figure S13.** Optimization of the dosage for a synthetic miR-631 mimic in SCC131 cells. Note, all the three dosages of the synthetic miR-631 mimic reduce the levels of both RAB11A transcript and protein. **, $p< 0.01$, ***,$p<0.001$. For the qRT-PCR, each bar is an average of two replicates (n=2). For the Western blot, both samples are processed in parallel and full uncropped blot images are given in supplementary Fig. S18. *Abbreviations*: MC, mimic control; and, nM, nano molar

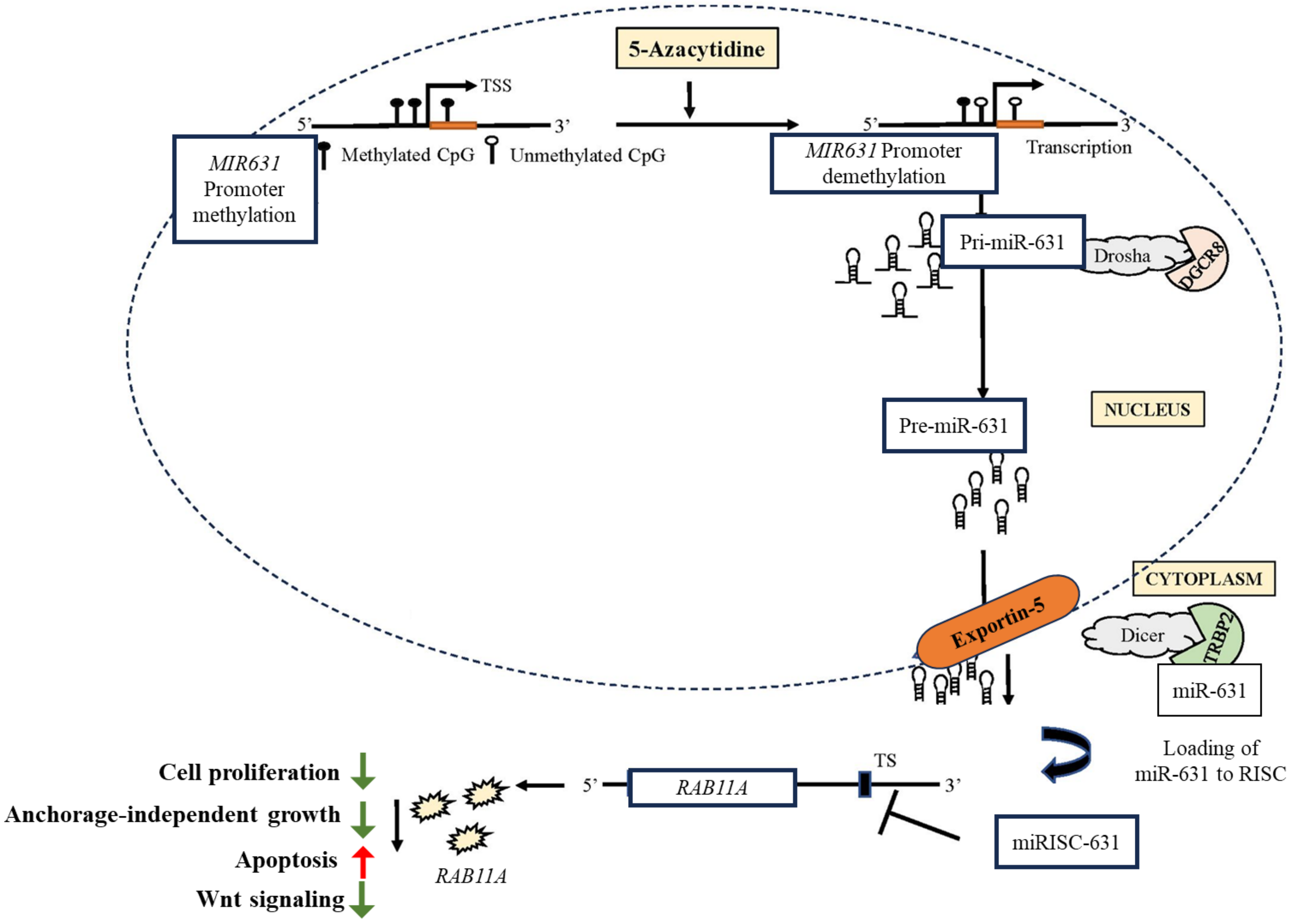


**Supplementary Figure S14. Summary of the study.** Diagrammatic summary of the regulation of the *MIR631* gene by promoter hypermethylation, biogenesis of miR-631, and the downstream functions of miR-631 in OSCC. *Abbreviations*: TSS, transcription start site; and, miRISC-631, microRNA induced silencing complex of miR-631. This figure is adapted from Kaushik et al., 2024 [15].
